# A novel vaccine strategy using quick and easy conversion of bacterial pathogens to unnatural amino acid-auxotrophic suicide derivatives

**DOI:** 10.1101/2023.09.14.557119

**Authors:** Yuya Nagasawa, Momoko Nakayama, Yusuke Kato, Yohsuke Ogawa, Swarmistha Devi Aribam, Yusaku Tsugami, Taketoshi Iwata, Osamu Mikami, Aoi Sugiyama, Megumi Onishi, Tomohito Hayashi, Masahiro Eguchi

**Affiliations:** National Institute of Animal Health, National Agriculture and Food Research Organization (NARO), Sapporo, Hokkaido, Japan; National Institute of Animal Health, National Agriculture and Food Research Organization (NARO), Tsukuba, Ibaraki, Japan; Institute of Agrobiological Sciences, National Agriculture and Food Research Organization (NARO), Tsukuba, Ibaraki, Japan

## Abstract

We propose a novel strategy for quick and easy preparation of suicide live vaccine candidates against bacterial pathogens. This method requires only the transformation of one or more plasmids carrying genes encoding for two types of biological devices, an unnatural amino acid (uAA) incorporation system and toxin-antitoxin systems in which translation of the antitoxins requires the uAA incorporation. *Escherichia coli* BL21-AI laboratory strains carrying the plasmids were viable in the presence of the uAA, whereas the free toxins killed these strains after removal of the uAA. The survival time after uAA removal could be controlled by the choice of uAA incorporation system and toxin-antitoxin systems. Multilayered toxin-antitoxin systems suppressed escape frequency to less than 1 escape per 10^9^ generations in the best case. This conditional suicide system also worked in *Salmonella enterica* and *E. coli* clinical isolates. The *S. enterica* vaccine strains were attenuated with a >10^5^-fold lethal dose. Serum IgG response and protection against the parental pathogenic strain were confirmed. In addition, the live *E. coli* vaccine strain was significantly more immunogenic and provided greater protection than a formalin-inactivated vaccine. The live *E. coli* vaccine was not detected after inoculation, presumably because the uAA is not present in the host animals or in the natural environment. These results suggest that this strategy provides a novel way to rapidly produce safe and highly immunogenic live bacterial vaccine candidates.

**Significance:** Live vaccines are the oldest vaccines with a history of more than 200 years. Due to their strong immunogenicity, live vaccines are still an important category of vaccines today. However, the development of live vaccines has been challenging due to the difficulties in achieving a balance between safety and immunogenicity. In recent decades, the frequent emergence of various new and old pathogens at risk of causing pandemics has highlighted the need for rapid vaccine development processes. We have pioneered the use of unnatural amino acids to control gene expression and to conditionally kill host bacteria as a biological containment system. This report highlights a quick and easy conversion of bacterial pathogens into live vaccine candidates using this containment system.

## Introduction

Emerging and reemerging infectious diseases are among the most serious threats to public health (1). High population densities due to urban concentration and increased contact between people around the world due to widespread air travel have greatly facilitated the global spread of pathogens (2,3). The emergence of pandemics in recent decades has highlighted the urgent need to develop strategies to control these infectious diseases (4). Vaccination is a promising strategy. Currently, the average development time of conventional vaccines from the preclinical stage is more than 10 years (5). However, the ongoing story of the fight against the global COVID-19 pandemic indicates the need for a much more rapid vaccine development strategy (6). Bacterial pathogens, including the ever-emerging antibiotic-resistant bacteria, pose a threat, as do viruses, including SARS coronaviruses, dengue viruses, and pandemic influenza viruses (7).

Today, new vaccine technologies such as virus-like particle vaccines, nanoparticle vaccines, DNA/RNA vaccines, and rational vaccine design are being developed in addition to traditional inactivated vaccines, live attenuated vaccines, viral vector vaccines, and subunit vaccines (8,9). Among these, live attenuated bacterial vaccines are the oldest in use and still one of the leading vaccine categories (10,11). Because live vaccines mimic natural infection almost perfectly, they can induce a broader range of immune responses in both humoral and cellular immunity. However, the development of live vaccines presents a unique dilemma (12). Namely, inadequate attenuation would result in high virulence and compromised safety, while excessive attenuation would compromise immunogenicity. This trade-off makes it difficult to obtain a live vaccine strain that balances immunogenicity and safety.

Here, we propose a quick and easy method to generate live vaccine candidates with balanced safety and immunogenicity using a conditional suicide system that can be achieved “additively” by plasmid transformation alone (Figure 1). This method is an application of our previously reported technique (13). This technique renders bacteria auxotrophic for an unnatural amino acid (uAA) that does not exist in the natural environment or in living organisms. Briefly, one or more toxin-antitoxin gene pairs are transformed into the pathogens using plasmids. We selected type II, type IV or type V toxin-antitoxins whose antitoxin genes encode proteins (14). A UAG stop codon(s) is inserted next to the translation initiation codon of the antitoxin genes. To incorporate the uAA at the UAG stop codon, the uAA-specific aminoacyl-tRNA synthetase and its cognate tRNA_CUA_ gene are also transformed (15). In the absence of uAA, no functional antitoxin is produced due to translation termination at the inserted UAG stop codon, resulting in killing of the host bacteria by the constitutively expressed toxin (16,17). In contrast, bacteria survive and proliferate in the presence of uAA because the toxin is neutralized by the functional antitoxin produced by the readthrough of the UAG stop codon. Thus, the pathogen can be converted into a conditional suicide strain that can only survive in the presence of the uAA by simply transforming these plasmids. For use as a live vaccine, the uAA-auxotrophic strains are first cultured in vitro in the presence of uAA to obtain the desired dose and then administered to humans or animals. Since the uAA have accumulated in the vaccine cells just after the supply is cut off, they survive for a while and could be immunogenic like the parental pathogens (13). Thereafter, the vaccine strain is expected to die over time and not cause disease.

**Figure 1.**
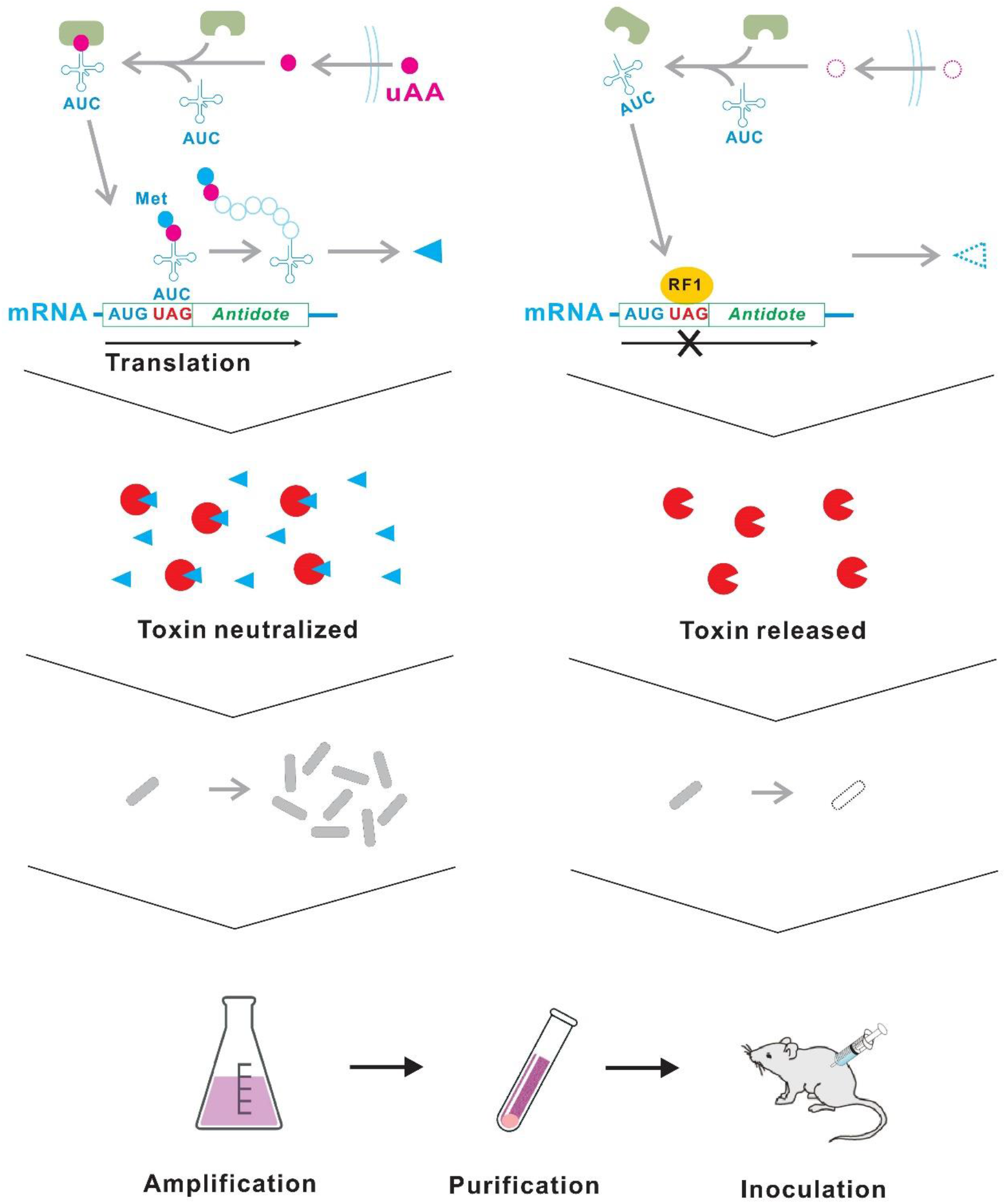
Schematic diagram of the uAA-auxotrophic suicide bacterial vaccine. Vaccine strains can grow in the presence of uAA. After purification from the culture medium, the bacterial vaccines are inoculated into humans or animals. After administration, the bacterial vaccine gradually dies due to depletion of the intracellular uAA. RF1, peptide release factor 1.

Unlike conventional live-attenuated vaccines, the uAA-auxotrophic strains are “suicide vaccines” that ensure safety by conditional suicide of the inoculated bacteria (18,19). Our method is expected to have the following features. First, candidate vaccine strains can be generated extremely rapidly. In modern live vaccine development, the induction of random or targeted mutations in genes of virulence-related and/or metabolic pathways is mainly used to attenuate wild-type pathogens (20). In this well-known method, vaccine strains are generated by “subtracting” genetic elements from the pathogen. This requires modification of the native genetic information of the pathogen, which is time consuming and labor intensive. Our method is extremely quick and easy because it is an “additive” method that only transforms the plasmids. This simple method should facilitate screening to select live vaccine strains with excellent safety and immunogenicity. Second, the attenuation can be controlled by the user, allowing a balance between safety and immunogenicity (12,21). Since the native genetic information of the parental pathogen is not altered, the immunogenicity should be maintained as original during the viability of the vaccine candidates. The duration of viability for the vaccine candidates would depend on the choice of toxin-antitoxins, uAA incorporation system and uAAs, suggesting that users can adjust the balance by adjusting these factors. Third, environmental risks are minimized. Live vaccines must be prevented from infecting unvaccinated individuals or spreading into the environment because of problems such as reversion to virulence, high susceptibility of immunocompromised individuals, and unintended immunization (22,23). Biological containment, in which the target organism is genetically programmed to survive only under the control of the users and to die in the natural environment, is desired to prevent the spread of live vaccines used in the open environment (24–27). Many biological containments using auxotrophy for natural nutrients or conditional toxin gene expression regulated by natural substances have often failed due to the presence of unexpected resources such as secretions and cell corpses (28,29). Therefore, auxotrophy for substances not present in the natural environment is an ideal biological containment (13, 30). In our method, live vaccine candidates must be supplied with a uAA in order to survive. The uAAs are not present in the natural environment or in living organisms, indicating that a high level of environmental safety should be ensured.

To demonstrate our idea, we constructed the suicidal *E. coli* laboratory strain BL21-AI auxotrophic for a uAA using transformation of the conditionally toxic plasmids. Various combinations of the uAA incorporation systems and the toxin-antitoxin systems were examined to achieve the requirements necessary for vaccine construction, including low escape frequency and control of survival time after uAA removal. In addition, the resulting conditional suicide system was applied to two species of bacterial pathogens, *S. enterica* and *E. coli* clinical isolates, to evaluate vaccine safety and immunogenicity. *S. enterica* is an intracellular parasitic human and zoonotic pathogen (31). *E. coli* is an important intestinal commensal bacterium of vertebrates, including strains that cause a variety of gastrointestinal and extraintestinal diseases in humans and livestock (32). Vaccination is a promising way to prevent both *S. enterica* and *E. coli* infections (33,34). Because *S. enterica* and *E. coli* are closely related phylogenetically (35), several plasmids replicate in both bacteria, including pBR322 and pACYC184, which harbor the pMB1 and p15A origin of replication, respectively (36). Thus, the *E. coli* BL21-AI conditional suicide systems were transferred intact to both *S. enterica* and *E. coli* clinical isolates. The resulting strains were inoculated into mice for vaccine evaluation.

## Results

### Development of uAA-regulated conditionally toxic plasmids

We first developed uAA-regulated conditionally toxic plasmids using the *E. coli* laboratory strain BL21-AI (37). Previously, we constructed the BL21-AI derivative, BL21-AI(IY-o), auxotrophic for the uAA 3-iodo-_L_-tyrosine (IY) by plasmid transformation (13) (Figure 2). BL21-AI(IY-o) carries two plasmids, plasmid B-1 which contains the IY-incorporation system consisting of the IY-specific aminoacyl-tRNA synthetase and its cognate tRNA_CUA_, and plasmid C1 which contains the toxin-antitoxin gene pair *colE3-immE3* (38) (Figure 2 and S1). The *immE3* contains a UAG-stop codon next to the translation start codon. BL21-AI(IY-o) died in the absence of IY, but more than 10^3^ escapers/10^8^ generations were detected, indicating that the escape frequency is >1 × 10^-5^ (13). This escape frequency is too high compared to the present standard for environmental safeguard set by National Institutes of Health (NIH), <1 × 10^-8^ (39).

**Figure 2.**
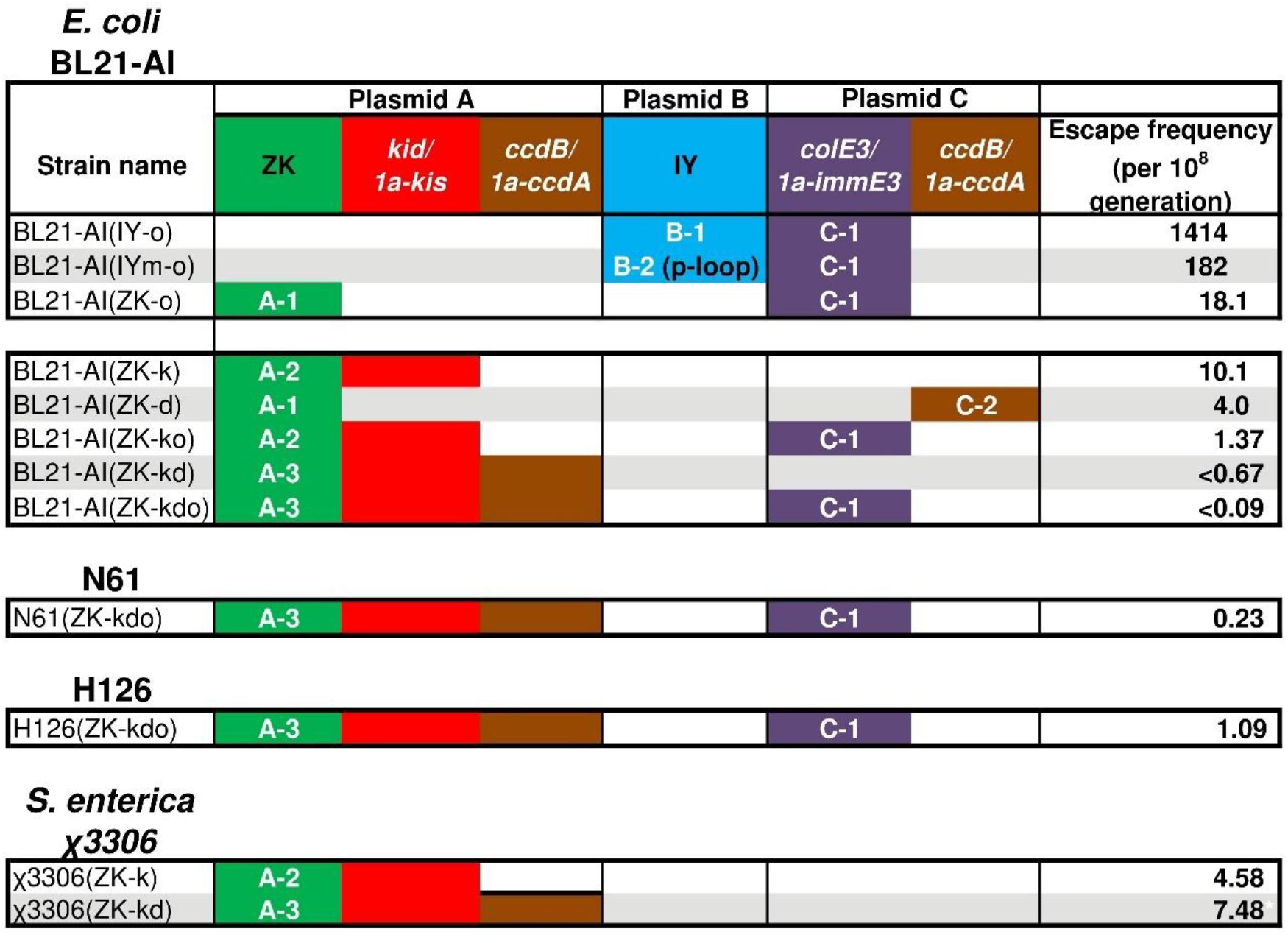
Escape frequency. Plasmids containing various uAA incorporation systems and toxin-antitoxin genes were transfected. Escape frequency in the absence of uAA was evaluated using a fluctuation assay. Values of the escape frequency are presented as escapers per 10^8^ generations. An inequality sign shows less than the indicated detection limit. ZK and IY, the ZK and IY incorporation system, respectively. A-1 to C-2, plasmid names shown in Figure S1.

A lower escape frequency reduces the risk of pathogenesis and increases the upper limit of safe dosage, as well as minimizing the environmental risk. Therefore, we modified the plasmids to reduce the escape frequency. Even in the absence of IY, the IY incorporation system translates UAG stop codon inserted mRNAs at 6-20% of the protein production in the presence of IY, presumably due to misincorporation of natural amino acids such as Tyr (16,40,41). The high-level leakage production of ImmE3 could reduce ColE3 toxicity and increase the escape frequency. A positive-feedback genetic circuit in plasmid B-2 suppresses the leakage production to <1% (42), resulting in a 0.1-fold reduction in escape frequency in BL21-AI(IYm-o) strain (Figure 2, S1 and S2). In addition, plasmid A-1, which contains a *N^ε^*-benzyloxycarbonyl-_L_-lysine (ZK) incorporation system consisting of the modified *Methanosarcina mazei* pyrrolysyl-tRNA synthetase specific for ZK and its cognate tRNA_CUA_ whose leakage production level is only 2%, was used in BL21-AI(ZK-o) instead of plasmid B-1 or B-2 (42,43). The escape frequency in BL21-AI(ZK-o) was further reduced to 0.1-fold of that in BL21-AI(IYm-o). Hence, we used the ZK incorporation system to construct strains with lower escape frequency.

Next, the effects of the toxin-antitoxin systems Kid-Kis and CcdB-CcdA were tested in addition to ColE3-ImmE3 (44,45). Instead of *colE3-immE3, kid-kis* and *ccdB-ccdA* were used in BL21-AI(ZK-k) and BL21-AI(ZK-d), respectively. Both BL21-AI(ZK-k) and BL21-AI(ZK-d) died in the absence of ZK, with an escape frequency comparable to that in BL21-AI(ZK-o) (Figure 2, S1 and S2). The escape frequency remained at >1 × 10^-8^.

Multi-layered containment systems, in which two or more kill switches are used together, have been reported to show a lower escape frequency (46,47). We, therefore, constructed strains carrying two or three toxin-antitoxin gene pairs. BL21-AI(ZK-ko) carrying both *colE3-immE3* and *kid-kis* showed a lower escape frequency than strains carrying either the toxin-antitoxin gene pair. The escape frequency in BL21-AI(ZK-kd) carrying both *kid-kis* and *ccdB-ccdA* reached at <0.67 × 10^-8^. Moreover, BL21-AI(ZK-kdo) carrying three toxin-antitoxin gene pairs, *kid-kis, ccdB-ccdA* and *colE3-immE3*, recorded the lowest escape frequency at <0.9 × 10^-9^. The escape frequencies in BL21-AI(ZK-kd) and BL21-AI(ZK-kdo) qualified the current NIH standard.

Changes in survival after the uAA removal were evaluated because the change may primarily affect how long the viable suicidal vaccine persists in the inoculated human/animal. BL21-AI(IY-o) continued to grow for at least 1 h after IY removal presumably due to intracellular accumulation, but then died rapidly with a half-life of approximately 50 min (13). It took around 6 h to decrease to 10% of the initial inoculation (T_0.1_). Growth rate may be another important factor determining the initial proliferation immediately after the uAA removal and the intracellular uAA consumption. BL21-AI(IY-o) grew at 0.63 times the rate of a vector control strain (13). Growth of BL21-AI(ZK-o) was only observed within 15-30 min, and T_0.1_ was 3-4 h (Figure 3A). In BL21-AI(ZK-kd), the growth period and T_0.1_ were largely extended to 2h and 4-5 h, respectively. Curiously, BL21-AI(ZK-kdo) had similar growth period and T_0.1_ to those of BL21-AI(ZK-o), 15-30 min and 2-2.5 h, respectively. The growth rates of BL21-AI(ZK-o), BL21-AI(ZK-kd) and BL21-AI(ZK-kdo) decreased to 0.89, 0.68 and 0.72 of the vector control strains, respectively, similar to BL21-AI(IY-o) (Table S1).

**Figure 3.**
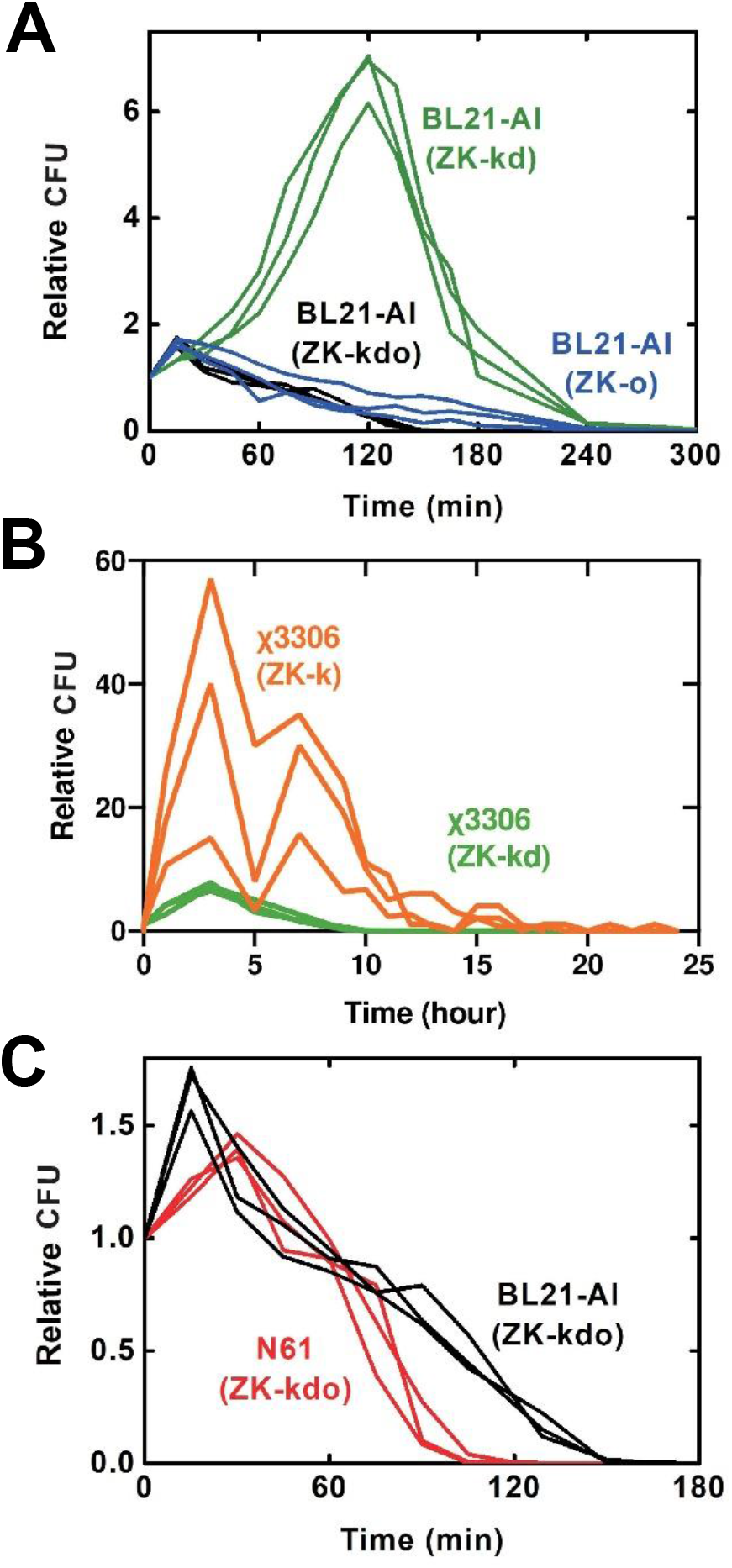
Viability after ZK removal. Some ZK-auxotrophic vaccine candidates were tested in the logarithmic growth phase. After exchange with ZK-free medium, viable bacterial cells were counted over time. Values are relative to T0. (A) BL21-AI strains carrying various combinations of toxin-antitoxins. (B) *S. enterica* χ3306(ZK-k) and χ3306 (ZK-kd) carrying A-2 and A-3, respectively. (C) Difference between BL21-AI and N-61 suicidal strains. Both strains carry A-3 and C-1 plasmids containing *kid-kis, ccdB-ccdA* and *colE3-immE3*.

### Application to S. enterica

We first constructed *S. enterica* suicide vaccine candidates using the uAA-regulated conditionally toxic plasmids. The plasmids developed using BL21-AI were applied for *S. enterica* without any modifications. A highly pathogenic strain against mice, χ3306, was used as a host (48). Plasmid A-2 or A-3 was successfully transformed into χ3306. The resulting strains were designated as χ3306(ZK-k) and χ3306(ZK-kd), respectively. Both χ3306(ZK-k) and χ3306 (ZK-kd) were ZK-auxotrophic (Figure S3). The escape frequency was slightly higher than the NIH standard (Figure 2). To further reduce the escape frequency, we attempted to transform plasmid C-1 into χ3306 (ZK-k) and χ3306 (ZK-kd), but no transformants were obtained. Changes in survival after the uAA removal for χ3306 (ZK-k) and χ3306(ZK-kd) were evaluated. The survival period of χ3306 (ZK-k) and χ3306(ZK-kd) were longer than those of BL21-AI strains (Figure 3B). In χ3306 (ZK-k), the growth periods and T_0.1_ were largely prolonged to 3-7h and 19-24h, respectively. Growth periods and T_0.1_ of χ3306 (ZK-kd) were 3h and 10h, respectively. Shape of the survival curve of χ3306 (ZK-kd), however, was similar to that of BL21-AI(ZK-kd). The growth rate of χ3306(ZK-k) and χ3306(ZK-kd) decreased to 0.62 and 0.93 of the vector control strains, respectively (Table S1).

To test the vaccine safety, χ3306 (ZK-k) and χ3306 (ZK-kd) were administered intravenously to BALB/c mice at 1 × 10^5^ CFU/mouse. Although all mice inoculated with the wild-type χ3306 were dead in 4 days after inoculation, all mice administrated with χ3306 (ZK-k) or χ3306 (ZK-kd) survived (Figure 4A). In addition, mice survived after intravenous administration of χ3306 (ZK-k) or χ3306 (ZK-kd) at 1 × 10^6^ CFU/mouse.

**Figure 4.**
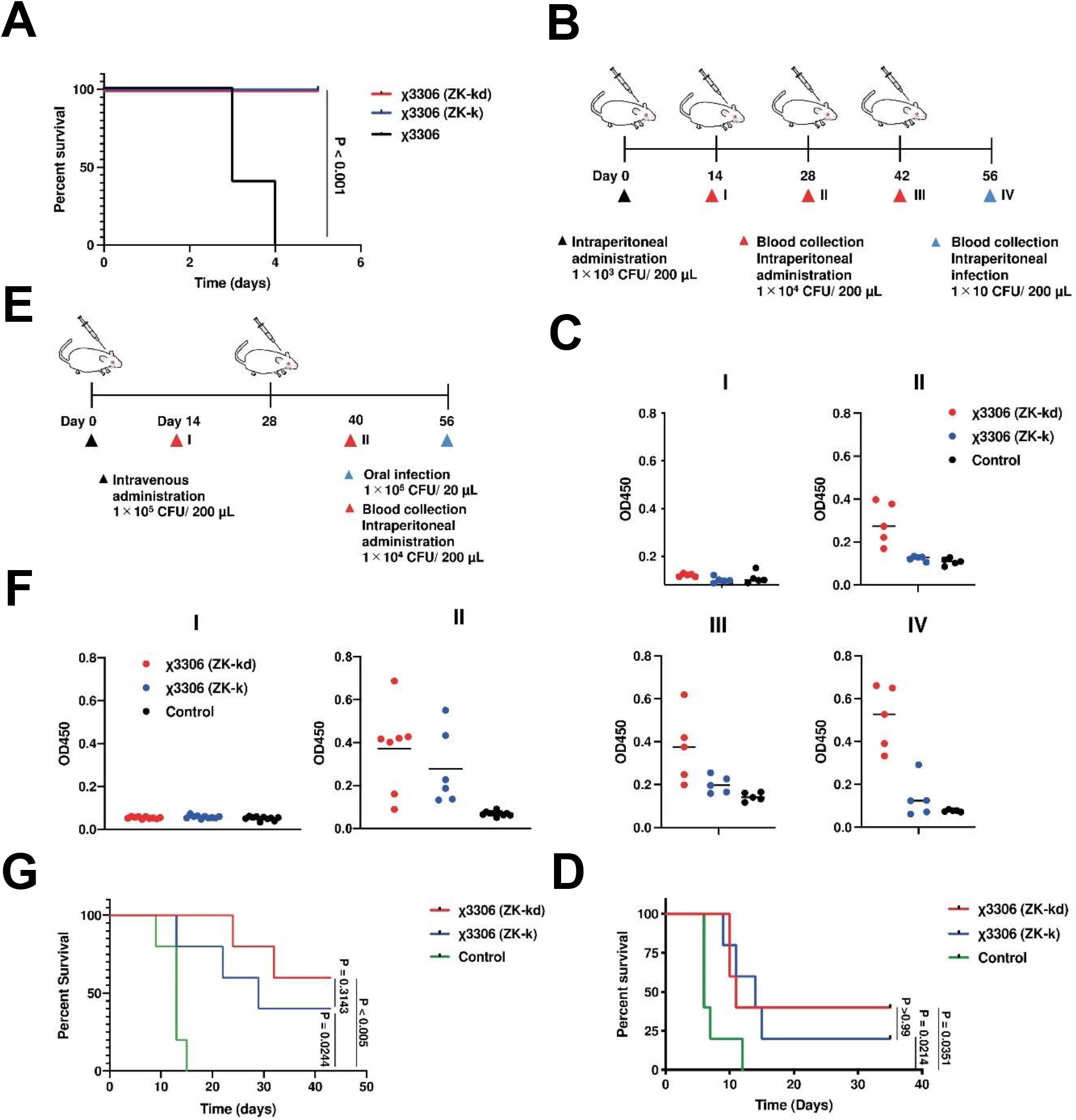
Application to *S. enterica*. (A) Survival curves of mice intravenously injected with *Salmonella* χ3306 (ZK-k) and χ3306 (ZK-kd), or wild-type χ3306 at the dose of 1 × 10^5^ CFU/mouse (n = 5). (B) Experimental design to assess the systemic humoral response and protection against *Salmonella* infection. BALB/c mice were immunized intraperitoneally with χ3306 (ZK-k) and χ3306 (ZK-kd). After 56 days, BALB/c mice were intraperitoneally infected. (C) Antibody titers of total IgG against LPS. I, II, III and IV indicate day 14, 28, 42 and 56, respectively. (D) Survival curves of *Salmonella* χ3306 (ZK-k) and χ3306 (ZK-kd) immunized mice intraperitoneally infected with χ3306 (wild-type) (n = 5). (E) Experimental design to assess protection against *Salmonella* oral infection by χ3306 (ZK-k) and χ3306 (ZK-kd) immunized mice. BALB/c mice were immunized intravenously with χ3306(ZK-k) and χ3306(ZK-kd), and 56 days later, BALB/c mice were orally infected. (F) Total IgG antibody titers against LPS. I and II indicate day 14 and 40, respectively. (G) Survival curves of *Salmonella* χ3306 (ZK-k) and χ3306 (ZK-kd) immunized mice orally infected with χ3306 (wild-type) (n = 5).

After fifth-dose vaccination, double-dose administration at 1 × 10^4^ CFU/mouse following triple-dose administration at 1 × 10^3^ CFU/mouse for χ3306 (ZK-k) or χ3306 (ZK-kd), the immunized mice were challenged with intraperitoneal injection of the wild-type χ3306 at 1 × 10 CFU/mouse (Figure 4B). We evaluated the induction of anti-*S. enterica* lipopolysaccharide (LPS) IgG after intraperitoneal administration with χ3306 (ZK-k) and χ3306 (ZK-kd) (Figure 4C). Significant induction of the anti-*S. enterica* LPS IgG was observed after double-dose vaccination with χ3306 (ZK-kd) at 1 × 10^3^ CFU/mouse or after double-dose vaccination of χ3306 (ZK-k) with an additional administration at 1 × 10^4^ CFU/mouse. χ3306 (ZK-kd) induced significantly more IgG than χ3306 (ZK-k).

Both χ3306 (ZK-k) and χ3306 (ZK-kd) vaccinated mice had 20% and 40% survival, respectively, at 35 days after the challenge. In contrast, all unvaccinated mice died by 12 days, suggesting that the vaccination with χ3306 (ZK-k) and χ3306 (ZK-kd) partially protected the mice against the wild-type χ3306 challenge (Figure 4D).

To determine the efficacy of protection against oral infections, we also performed an oral challenge test (Figure 4E). Mice were intraperitoneally vaccinated with a double-dose administration at 1 × 10^5^ CFU/mouse of χ3306 (ZK-k) or χ3306 (ZK-kd). When total IgG titers were assayed, production was observed after two inoculations (Figure 4F). Immunized mice were challenged orally with 1.5 × 10^5^ CFU/mouse of wild-type χ3306. All non-immunized mice died until 15 days after the challenge. Mice immunized with χ3306 (ZK-k) or χ3306 (ZK-kd) survived for the 42 days (60% and 40%, respectively) (Figure 4G).

Conclusively, the ZK-auxotrophic suicidal *S. enterica* vaccine is less virulent and immunogenic enough to induce antibody production and confer protection against the wild-type parent strain. Therefore, the highly virulent strain χ3306 has been successfully transformed into a live vaccine.

### Application to E. coli clinical isolates

Following the promising results with *S. enterica*, the ZK-auxotrophic suicide vaccine was subsequently constructed in clinical isolates of *E. coli.* Six non-enterohemorrhagic *E. coli* strains, H19, H20, H126, N61, N67 and N77 were isolated from the milk of cows with coliform mastitis (49) and were subjected to the plasmid transformation.

First, we tested the tolerance for ZK incorporation at the UAG stop codons. All tested clinical isolates carrying plasmid A-1, which contains only the ZK incorporation system and no toxin-antitoxin genes, were viable in the presence of ZK (Figure S1 and S4), suggesting that all tested strains are tolerant to ZK incorporation.

Next, plasmid A-3 containing the ZK incorporation system, *kid-kis* and *ccdB-ccdA* was transformed into the parental clinical isolates. Some strains carrying plasmid A-3, H19((ZK-kd), H126(ZK-kd), N61(ZK-kd) and N67(ZK-kd), were successfully isolated although others, H20 and N77, could not be isolated even after several attempts (Figure S5). No survivors were detected in H126(ZK-kd), N61(ZK-kd) and N67(ZK-kd) after inoculation with 5 × 10^4^ CFU in the absence of ZK. Nonetheless, several tens of survivors were observed in H19(ZK-kd), which was excluded from further plasmid transformation.

In addition, plasmid C-1 containing *colE3-immE3* was transformed into H126(ZK-kd), N61(ZK-kd) and N67(ZK-kd). Bacteria carrying both plasmids A-3 and C-1 were successfully isolated for H126 and N61, while N67 was excluded due to natural resistance to the selection antibiotic of plasmid C-1, carbenicillin (Figure S6).

Finally, we obtained two strains carrying the ZK-incorporation system with 3 inducible toxin-antitoxin pairs, N61(ZK-kdo) and H126(ZK-kdo). The escape frequencies were 0.23 and 1.09 per 10^8^ generation for N61(ZK-kdo) and H126(ZK-kdo), respectively (Figure 2). Safety and immunogenicity was verified using N61(ZK-kdo) which satisfied the NIH standard for escape frequency. N61(ZK-kdo) was killed slightly faster after ZK-removal than BL21-AI(ZK-kdo) (Figure 3C). The T_0.1_ of N61(ZK-kdo) was estimated to be 90-105 min. Growth rates of N61(ZK-kdo) decreased to 0.65-fold that of the N61 vector control strain.

To confirm the safety of the N61(ZK-kdo) vaccine, we evaluated the safety profiles using a single-dose vaccination schedule in BALB/c mice with the N61(ZK-kdo), formalin-killed N61(ZK-kdo) or wild-type N61 at 1 × 10^8^ CFU (Figure 5A). In the treatment of wild-type N61 by subcutaneous injection, all mice were developed terminal disease and died. In comparison, all mice were survived in treatment with N61(ZK-kdo), formalin-killed N61(ZK-kdo) and saline. Clinical signs in all vaccinated mice started with ruffled fur, hunched posture, decreased activity, and progressed to limb paresis or paralysis, tremor, ataxia, rigidity, dehydration, and coma occurring within 12 to 48 hours of challenge, and then surviving mice recovered within 60 hours (Figure 5B, Table S2). Tissue specimens of wild-type N61 at 24 hours after injection showed bacterial colonization that was not seen in N61(ZK-kdo) and saline injection (Figure 5C). In addition, viable wild-type N61 bacteria were detected in the blood samples at 12 hours after injection but not at 3 hours, indicating bacterial proliferation after the injection (Figure 5D). In contrast, we could not detect any viable bacteria in the samples obtained from mice injected with N61(ZK-kdo).

**Figure 5.**
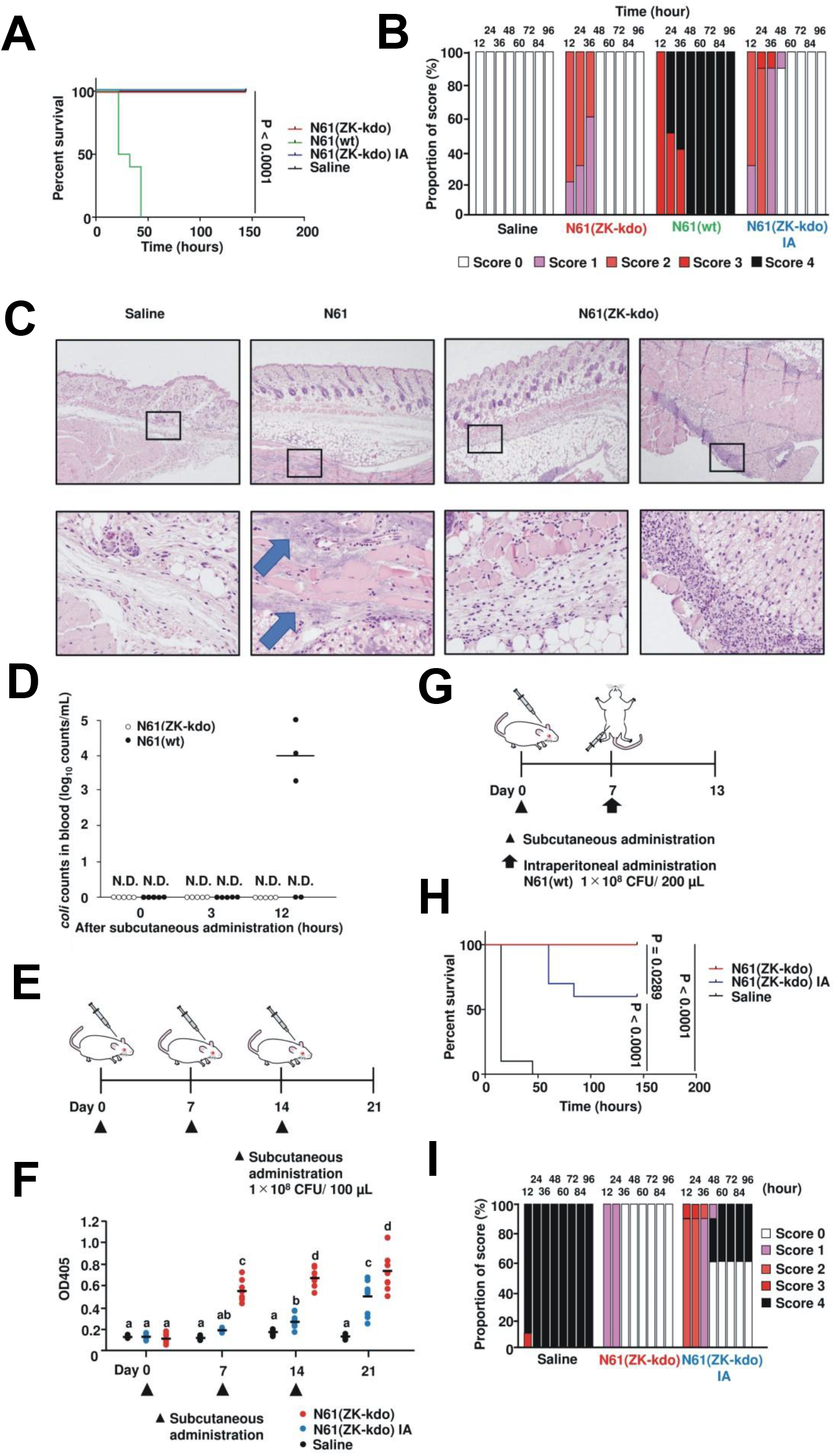
Application to *E. coli* clinical isolates. (A) Survival curve of mice after subcutaneous administration of N61, N61(ZK-kdo) or formalin-killed N61(ZK-kdo) strain at 1 × 10^8^ CFU during the course of 144 hours period (n = 10). (B) Clinical scoring results over the course of sepsis in mice that subcutaneously administrated N61, N61(ZK-kdo) or formalin-killed N61(ZK-kdo) strain. The scoring criteria are presented in Table S2. Bars show the distribution of scores for the observation parameters calculated based on scoring 1, 2, 3, and 4 at the time points after subcutaneous administration. The scores are represented by distinct colors as indicated. (C) Histological specimens of subcutaneous mouse tissues harvested at 24 hours. Square: large scale of bottom figures. Arrows: bacterial aggregation. (D) Kinetics of blood clearance showing the complete elimination of N61(ZK-kdo) after subcutaneously administration in mice. White and black circles correspond to blood samples from N61(ZK-kdo) and wild-type N61 strain subcutaneously administrated mice, respectively. N.D., not detected. (E) Experimental design for evaluation of the systemic humoral responses in mice by subcutaneous administration of N61(ZK-kdo). Mice were immunized with N61(ZK-kdo) or formalin-killed N61(ZK-kdo) strain by subcutaneous administration. After three subcutaneous doses every week, anti-*E. coli*-specific IgG antibodies in serum were determined at each time point. Triangles: time point of subcutaneous administration. (F) Anti-*E. coli*-specific IgG titers in serum after subcutaneous administration of N61(ZK-kdo). Each data point represents the OD value for one sample. The bar represents the mean. Circles are represented by distinct colors as indicated. a-c: The values (average ± SD; n = 10) between different superscripts are significantly different (p < 0.05). (G) Experimental design for evaluation of the protection conferred by subcutaneous administration of N61(ZK-kdo). Mice were immunized with N61(ZK-kdo) or formalin-killed N61(ZK-kdo) strain by subcutaneous administration. After one-week subcutaneous doses, mice were infected with wild-type N61 strain (1 × 10^8^ CFU) by intraperitoneal administration. (H) Survival curves of mice administrated with N61 strain during 144 hours (n = 10). (I) Clinical scoring results over the course of sepsis in subcutaneously administrated mice that intraperitoneally administrated N61 strain.

To measure the antibody-mediated immune responses following vaccination with N61(ZK-kdo), BALB/c mice were immunized with N61(ZK-kdo), formalin-killed N61(ZK-kdo) or saline (Figure 5E and 5F). After a three-dose vaccination schedule, significant levels of total IgG against *E. coli* were present in all immunized mice on days 7 and 14 after vaccination with N61(ZK-kdo) compared to saline, whereas no significant increase was detected in mice immunized with formalin-killed N61(ZK-kdo). On day 21 after vaccination, total IgG levels increased significantly in both N61(ZK-kdo) and formalin-killed N61(ZK-kdo) immunized mice. Notably, the live N61(ZK-kdo) was more immunogenic than the formalin-killed form.

To verify whether N61 (ZK-kdo) confers protection, we assessed efficacy using an acute systemic infection model in mice (Figure 5G-5I). A single-dose vaccination schedule with the N61 (ZK-kdo) protected mice against challenge with the wild-type N61 on day 7, whereas all control mice died within 48 h. Similarly, the survival rate of vaccinated mice with formalin-killed N61 (ZK-kdo) was 60%. In contrast, all vaccinated mice with N61(ZK-kdo) were alive. Clinical signs in all mice started with ruffled fur, hunched posture, decreased activity, and progressed to limb paresis or paralysis, tremor, ataxia, rigidity, dehydration, and coma occurring within 12 to 48 hours after challenge. The surviving mice recovered within 60 hours.

## Discussion

We have demonstrated a novel platform for quick and easy preparation of suicide vaccine candidates in *S. enterica* and *E. coli*. This platform is a transformation-based system using plasmids containing a uAA incorporation system and a toxin-antitoxin system(s) regulated by an externally supplied uAA.

The selection of the uAA incorporation system affected escape frequency. In the IY incorporation system, a modified system with better switchability using a positive-feedback loop reduced the escape frequency to 0.1-fold. The modified system suppresses leakage expression, indicating that lowered antitoxin expression level in the absence of IY enhances killing by the cognate toxin. Exchange the modified IY incorporation system with the ZK incorporation system further reduced the escape frequency to 0.1-fold. In addition, both the initial proliferation period and duration of viable bacteria after the uAA removal were also reduced. Since the switchability of the modified IY and ZK incorporation systems is at a similar level, a difference in biological or chemical properties between IY and ZK, or a lower efficiency of the ZK incorporation is a possible cause, which could affect the stability, activity or expression level of the antitoxins.

Incorporation of uAA sometimes severely damages host bacteria (50). However, all the tested strains, including the *E. coli* laboratory strain BL21-AI, *E. coli* clinical isolates and *S. enterica* χ3306, were tolerant to the incorporation of ZK at the UAG stop codon. This observation suggests that the disorder caused by the uAA incorporation may not be a frequently occurring barrier whereas more extensive research would be needed.

Selection and combination of the toxin-antitoxin systems was crucial in determining the vaccine characteristics. We showed that, in addition to previously used ColE3-ImmE3, toxicity of Kid-Kis and CcdB-CcdA can be controlled by ZK in *E. coli* BL21-AI, suggesting that the repertoire of available toxin-antitoxins is scalable. However, plasmid A3 containing both *kid-kis* and *ccdB-ccdA* failed to be transformed into the *E. coli* clinical isolates H20 and N77, suggesting that the toxicity was not completely suppressed in these strains even in the presence of ZK. Similarly, *S. enterica* χ3306 was unable to maintain plasmid C-1 containing *colE3-immE3*. Conversely, unusually many survivors were detected in H19 carrying plasmid A3, presumably due to lower toxin or predominant antitoxin activity. These observations suggest that the toxin-antitoxin systems working well in BL21-AI are not always available in other *E. coli* and *S. enterica* strains. The gene organization including regulatory regions was maintained as wild-type in the toxin-antitoxin genes because the expression intensity of these genes is optimally balanced by evolutionary selection for stable maintenance through generations (13,51). The optimal gene organization in one strain, nonetheless, may not necessarily be the case in other strains. Moreover, the antitoxin protein production could be attenuated due to competition between ZK-tRNA_CUA_ and peptide release factor 1 at the UAG stop codons (52,53), thereby disturbing the optimal balance between toxins and antitoxins. A possible solution is to use a wide variety of toxin-antitoxin genes and a library of mutants of their regulatory regions. The most promising vaccine candidates with desired characteristics may be obtained after screening of the library transformants.

All strains carrying only a single toxin-antitoxin were unable to reach the escape frequencies below 1 × 10^-8^. Some toxin-antitoxins, nonetheless, were available in this study, enabling to construct multilayered systems. Finally, we achieved to generate strains with low escape frequencies below 1 × 10^-8^ using a combination of two or three toxin-antitoxins, suggesting that the multilayered system is an effective strategy also in our method, as suggested in previous studies on biological containment systems (46,47).

The duration of bacterial viability after ZK removal was affected by selection of toxin-antitoxins. In BL21-AI, the strain carrying both *kid-kis* and *ccdB-ccdA* survived for relatively long time, but additional transformation of *colE3-immE3* dramatically reduced the duration. Interestingly, the reduced duration was almost identical to that of *colE3-immE3* single transformant. In this case, the duration should be determined by the toxicity of the specific toxin-antitoxin rather than the sum of the effects of multiple toxin-antitoxins since a single toxin is enough to kill the host. This observation is of great significance in selecting the appropriate toxin-antitoxin combination to achieve the intended duration. χ3306(ZK-kd) continued to proliferate and survived remarkably longer after ZK removal than BL21-AI(ZK-kd), suggesting that host-dependent differential responses are another important factor.

All the vaccine candidate strains showed impaired growth rates compared to the vector control. This secondary attenuation should be considered as a factor affecting vaccine efficacy.

Safety is one of the most important characteristics since we intend to convert pathogens directly into vaccine candidates. Mice survived after inoculation with *S. enterica* χ3306(ZK-k) and χ3306(ZK-kd) at 1 × 10^5^ CFU although the LD_50_ value of wild-type χ3306 is <1 × 10 CFU. Wild-type *E. coli* N61 killed mice at 1 × 10^8^ CFU and formed bacterial colonies near the injection site. In addition, the viable bacteria were detected in blood 12 h after injection indicating the bacterial proliferation *in vivo*. In contrast, mice were survived after the injection of N61(ZK-kdo) at the same dose. No colony formation or proliferation was observed *in vivo*. These results suggest that the ZK-auxotrophic suicidal vaccine candidates are much less virulent than the parental wild-type pathogens in both *S. enterica* and *E. coli* clinical isolates.

Another critical characteristic is immunogenicity. Specific IgG induction and protection against the parental wild-type pathogens were confirmed in both *S. enterica* and *E. coli.* Two important observations were noted. First, protection against oral challenge, as well as intraperitoneal challenge, was observed in *S. enterica*. Second, the *E. coli* suicide live vaccine induced stronger IgG production and showed more potent protection than the formalin-inactivated vaccine. These findings suggest that the strong immunogenicity characteristic of excellent live vaccines is achievable in the uAA-regulated conditional suicide vaccines.

We selected two bacterial pathogens, *S. enterica* and *E. coli*, as models for vaccine construction. *S. enterica* encompasses *S. typhi* and *S. paratyphi,* which respectively cause typhi and paratyphi fever (54). Non-typhoidal salmonella is also a major public health problem in Africa, killing ∼49,600 people each year (55). *S. enterica* is pathogenic in animals as well as humans. Domestic salmonellosis has been reported in cattle, swine, and chickens. Most cases of salmonellosis are caused by the consumption of contaminated eggs, chicken, pork, beef, and dairy products, although other animals such as rats and companion animals have also been identified as sources of infection in human. *S. enterica* has the potential to be drug-resistant, and vaccination is an effective measure for controlling infections. However, no universal vaccine is currently available for humans and animals. Animal studies have shown that attenuated live vaccines are more effective, but they may pose a danger to immunodeficiency patients and children. χ3306(ZK-k) and χ3306(ZK-kd)-immunized mice showed protection against infection both peritoneally and orally. Furthermore, these vaccine strains were dramatically less virulent than wild-type χ3306. These results suggest that the uAA-auxotrophic suicide vaccine represents a promising approach for developing safe and highly immunogenic *Salmonella* vaccines.

Pathogenic *E. coli,* which has acquired specific combinations of virulence factors, causes various diseases in healthy individuals, including enteric/diarrheal disease, urinary tract infections and sepsis/meningitis (32). *E. coli* is also an important pathogen in animals. *E. coli* mastitis is one of the most common diseases in the dairy industry and can cause severe acute inflammation with fever and decreased milk production, and lead to serious economic losses (56–59). The virulence factors of the pathogenic *E. coli,* which include LPS among others, have been documented (60). During *E. coli* infection and multiplication in the mammary gland, the release of LPS activates the host’s immune system (61). Although immune cells can invade and destroy LPS, the large amount of LPS is released and causes systemic symptoms in the host (62). Currently, the vaccine used to prevent *E. coli* bovine mastitis is the J5 vaccine which is an inactivated *E. coli* vaccine with an incomplete O-antigen (62). A previous study showed that the J5 immune serum was not an improvement on the already high efficiency of naturally acquired antibodies against *E. coli* (63). Moreover, the safety and effectiveness of the J5 vaccine in dairy cows has not been clearly reported (64). The use of killed *E. coli* vaccines including autovaccines has not been produced the desired effects (65,66). The attenuation procedure should be performed in a manner that prevents the vaccine from causing a persistent carrier state (67). This can be achieved by N61(ZK-kdo) strain’s ability to colonize and proliferate in the host for a limited period being eliminated without causing disease. This was observed for the N61(ZK-kdo) when injected intraperitoneally into mice, as it was completely cleared from the blood. N61(ZK-kdo) demonstrated superior vaccine efficacy compared to formalin-inactivated N61(ZK-kdo), suggesting that this innovative vaccine is a possible strategy to protect against various human and animal colibacilloses including bovine coliform mastitis.

In conclusion, we demonstrated the basic concept of a novel strategy for quick and easy conversion of bacterial pathogens into the uAA-auxotrophic suicide vaccine candidates using the conditionally toxic plasmids. Users can control the critical vaccine characteristics such as the escape frequency and the duration of bacterial viability after the uAA removal by selecting of the uAA incorporation system and the toxin-antitoxins. This platform should enable the rapid preparation of vaccine candidate strains against various infectious diseases caused by *S. enterica* and *E. coli* for which highly effective vaccines have not yet been developed. The concept of the uAA-auxotrophic suicide vaccine may be scalable for other microbial pathogens after transplanting the uAA incorporation systems and developing the applicable toxin-antitoxin systems.

## Materials and Methods

### Strains, growth conditions, and transformation

*E. coli* BL21-AI was commercially purchased. Six non-enterohemorrhagic *E. coli* strains, H19, H20, H126, N61, N67 and N77, were isolated from milk of cows with coliform mastitis in Hokkaido, Japan. *E. coli* XL1-Blue and *E. coli* DH5α were also used for plasmid construction. All *E. coli* strains were grown in Luria-Bertani (LB) medium or its dilutions at 37°C and 200 rpm for all experiments. *Salmonella enterica* serovar Typhimurium (*S.* Typhimurium) χ3306 was grown in LB medium at 37°C and standing culture for all experiments. Agar (final 2%) was added for preparation of solid medium. Carbenicillin (100 µg/ml) and chloramphenicol (50 µg/ml) were added as appropriate. Transformation was performed by electroporation using a Gene Pulser II^TM^ electroporator (BIO-RAD).

### Plasmid construction

Maps for all plasmids are shown in Figure S1. Plasmid A-1 and B-1 are identical to pTYR MjIYRS2-1(D286) MJR1 × 3 and pTK2-1 ZLysRS1, respectively, kindly provided from Kensaku Sakamoto and Shigeyuki Yokoyama (RIKEN) (41,43). Plasmid C-1 is pSH350 distributed by Haruhiko Masaki (38). Plasmid A-2, A-3 and C-2 were constructed using gene synthesis outsourced to GenScript (Tokyo, Japan). Plasmid B-2 is 2a-IYRS(M6V) described in our previous paper (42).

### Escape frequency

Escape frequency was estimated by fluctuation assay as previously described (13, 68), with some modifications. Briefly, ten parallel cultures were prepared using two-fold diluted LB medium containing 3 mM ZK for *E. coli* strains and 1mM ZK for *S.* Typhimurium strains and incubated for approximately 16 h (final OD_590_ = 0.03-0.1). The bacteria were collected by centrifugation and resuspended in 1 ml of ZK-free LB medium and centrifuged again for washing. The wash was repeated three times to remove ZK completely. The bacteria were resuspended and inoculated onto ZK-free solid medium. After overnight culture, escapers were detected as colonies. To estimate total number of inoculated bacteria, the bacterial suspension was also plated onto solid medium containing 3 mM ZK for *E. coli* strains and 1mM ZK for *S.* Typhimurium strains. The mutation rates were calculated by the web tool FALCOR using the Ma-Sandri-Sarkar Maximum Likelihood Estimator method (https://lianglab.brocku.ca/FALCOR/) (69).

### Duration of bacterial viability after ZK removal

Bacteria were cultured in LB medium diluted to 1/4 concentration containing 3 mM ZK for *E. coli* strains and 1mM ZK for *S.* Typhimurium strains and appropriate selection marker antibiotics for 16 h. To obtain the bacteria at the logarithmic growth phase, the culture medium was exchanged with fresh ZK-containing LB medium. The bacteria were additionally cultured for 1.5 h. After washing twice with fresh LB medium, the bacterial suspension was diluted 10^5^-fold in 30 ml of ZK-free LB medium and incubated. An aliquot (250 μl) of the bacterial culture was directly inoculated onto a ZK-containing LB agar plate. After overnight culture, appearing colonies were counted.

### Growth rate

The bacterial culture was prepared as described above and diluted in 50 ml of LB medium containing 3 mM ZK for *E. coli* strains and 1mM ZK for *S.* Typhimurium strains to calculated OD_590_ = 0.01-0.03. The diluted culture was incubated for 1-2 h until OD_590_ reached to 0.6-1.2. An aliquot (1 ml) of the bacterial culture was withdrawn at 15 min interval to measure bacterial density as OD_590_. A single fitted curve was generated for each strain using Origin7.

### Animal

Three-week-old female BALB/c mice were purchased from SLC Japan, Inc. (Hamamatsu, Japan) and Hokudo Co. (Sapporo, Japan). Animals were maintained and used in accordance with the Guidelines for the Care and Use of Laboratory Animals of the National Institute of Animal Health. All animal procedures were carried out in strict accordance with local guidelines and with ethical approval from the National Institute of Animal Health (authorization number: 19-035 and 20-081). Blood samples were collected from the submandibular vein of anesthetized mice, and sera were separated from the blood cells by centrifugation (1,500 g, 15 min) and stored at −80 °C until subsequent analysis.

### *Salmonella* Infection experiments

Mice were immunized by intraperitoneal and intravenous administration of χ3306 (ZK-k) or χ3306 (ZK-kd), all challenges were made using the intraperitoneal administration route and oral challenge with *S.* Typhimurium χ3306. Intraperitoneal infections were given at a dose of approximately 10 CFU/200 μl (LD_50_ in BALB/c: <10 CFU) (70,71). Oral infections were administered at approximately 10^6^ CFU/20 μl (LD_50_ in BALB/c: <10^4^ CFU) (72).

### *E. coil* Infection experiments

To prepare the formalin-inactivated *E. coli* N61 (ZK-kdo), the washed bacteria were resuspended in phosphate-buffered saline (PBS) containing 0.5% formaldehyde, incubated with shaking (50 rpm) at room temperature for inactivation, and then washed three times with PBS. After 1 h following inactivation, the cells were checked by plating on LB agar to determine the absence of colonies after overnight incubation. The inactivation step was then performed overnight to ensure complete inactivation.

All immunizations were made using the subcutaneous administration, all challenges were performed by the intraperitoneal administration route of administration. N61 and N61 (ZK-kdo) strains were cultured in LB with 10 mM ZK at 37 °C and 180 rpm, harvested (8,000 g, 15 min), washed twice with LB and finally adjusted with saline. For intraperitoneal and subcutaneous administration of N61 and N61 (ZX-kdo) strains, we used a total volume of 200 and 100 μl, respectively. During the subcutaneous administration or infection with *E. coli,* mice were monitored twice daily and a final disease score was given to each mouse according to the clinical signs observed as summarized in Table S2 (73).

### Histological analysis

Tissue samples collected from the subcutaneous administration site were fixed in 10% phosphate-buffered formalin and embedded in paraffin for histochemical analysis. Paraffin sections (4 μm thick) were mounted on slide glasses, de-waxed in xylene, rehydrated through a series of graded ethanol solutions, and transferred to PBS (pH 7.4). Tissue sections were stained with hematoxylin and eosin (HE). The sections were observed using a light microscope (Leica, Eclipse Ni-U, Wetzlar, Germany).

### Enzyme-linked immunosorbent assay (ELISA) for the detection of specific IgG antibodies

For ELISA, a 96-well plate (Nunc; Roskilde, Denmark) was coated with 50 μl of 10 μg/ml LPS from *S.* Typhimurium χ3306 diluted in 50mM carbonate-bicarbonate buffer, pH 9.6. All incubations were carried out at 37°C for 60 min, and every incubation step was followed by four washes with ELISA wash buffer (0.9% NaCl supplemented with 0.1% Tween-20). After coating and washing, the plates were incubated with serum samples diluted 1:100 in PBST (PBS with 0.1% Tween-20). Bound antibodies were detected by using goat anti-mouse IgG conjugated to horseradish peroxidase (HRP; Southern Biotech, Birmingham, AL, USA) diluted 1:2000 in PBST and developed using 3,3’,5,5’-tetramethylbenzidine substrate (Thermo Scientific Pierce, Rockland, IL, USA). Absorbance was measured at 450 nm using an ELISA plate reader.

To detect specific IgG antibodies against *E. coli* in serum, a microtiter plate was directly coated with formalin-killed wild-type N61 as capture antigen. Formalin-killed N61 in PBS was dried in an oven at 5×10^6^ cells/well in 96-well ELISA plates (C96 Maxisorpcert, Nunc-Immuno Plate, Thermo Fisher Scientific) overnight at 37°C. After incubation, wells were washed with Tris-buffered saline-Tween-20 (TBST) and then incubated with 100 μl of milk (diluted 1:100 in PBS) for 90 min at room temperature. After five TBST washes, wells were incubated with horseradish peroxidase-conjugated sheep anti-mouse IgG antibody (diluted 1:30,000, Bethyl Laboratories, Inc., Montgomery, TX, USA) for 2 h at room temperature. The freshly prepared substrate was added, and OD was measured at 405 nm using the 3,3’,5,5’-tetramethylbenzidine microwell peroxidase substrate system (KPL, Gaithersburg, MD, USA). All samples were analyzed in duplicate and mean values were calculated. Anti-E. coli-IgG antibodies were calculated by subtracting the OD values for the buffer controls (OD_405_ = ∼0.10), which were included in duplicate in all ELISAs, from the specific sample OD values. To standardize and compare results between plates, positive control milk samples (from murine serum IP by *E. coli*, OD_405_ = ∼1.0 for specific IgG) were included in duplicate in all ELISAs. OD values were normalized against those of positive controls.

### Statistical analysis

The one-way analysis of variance (ANOVA) was used for multiple comparisons, followed by Bonferroni’s post hoc test using SPSS software (IBM SPSS Statistics version 25, Tokyo, Japan). Survival data were compared using the log-rank test and Wilcoxon test using Prism 6.0 (GraphPad Prism, GraphPad Software, Inc.). Differences with P values less than 0.05 were considered significant.

### Data, Materials, and Software Availability

All data is included in the manuscript and/or supporting information.

## Acknowledgments

We thank the following researchers: Kensaku Sakamoto and Shigeyuki Yokoyama (RIKEN) for the Plasmid A-1 and B-1. Haruhiko Masaki (Tokyo University) for Plasmid C-1. This work was supported by JSPS grant number 20H03158.

## Author contributions

Y.K. designed the project; Y.K. designed microbial experiments; M.E. and M.N. designed *S. enterica* vaccine experiments; Y.N. and T.H. designed *E. coli* vaccine experiments; Y.N., M.N., Y.K., Y.O., S.D.A., Y.T., T.I., O.M., A.S., M.O., T.H., and M.E. performed research; Y.N., M.N., Y.K., T.H., and M.E. analyzed data; Y.K., Y.N., M.N., and M.E. wrote the paper. All authors reviewed drafts of the paper.

## Supplementary information

**Supplementary Figure 1.**
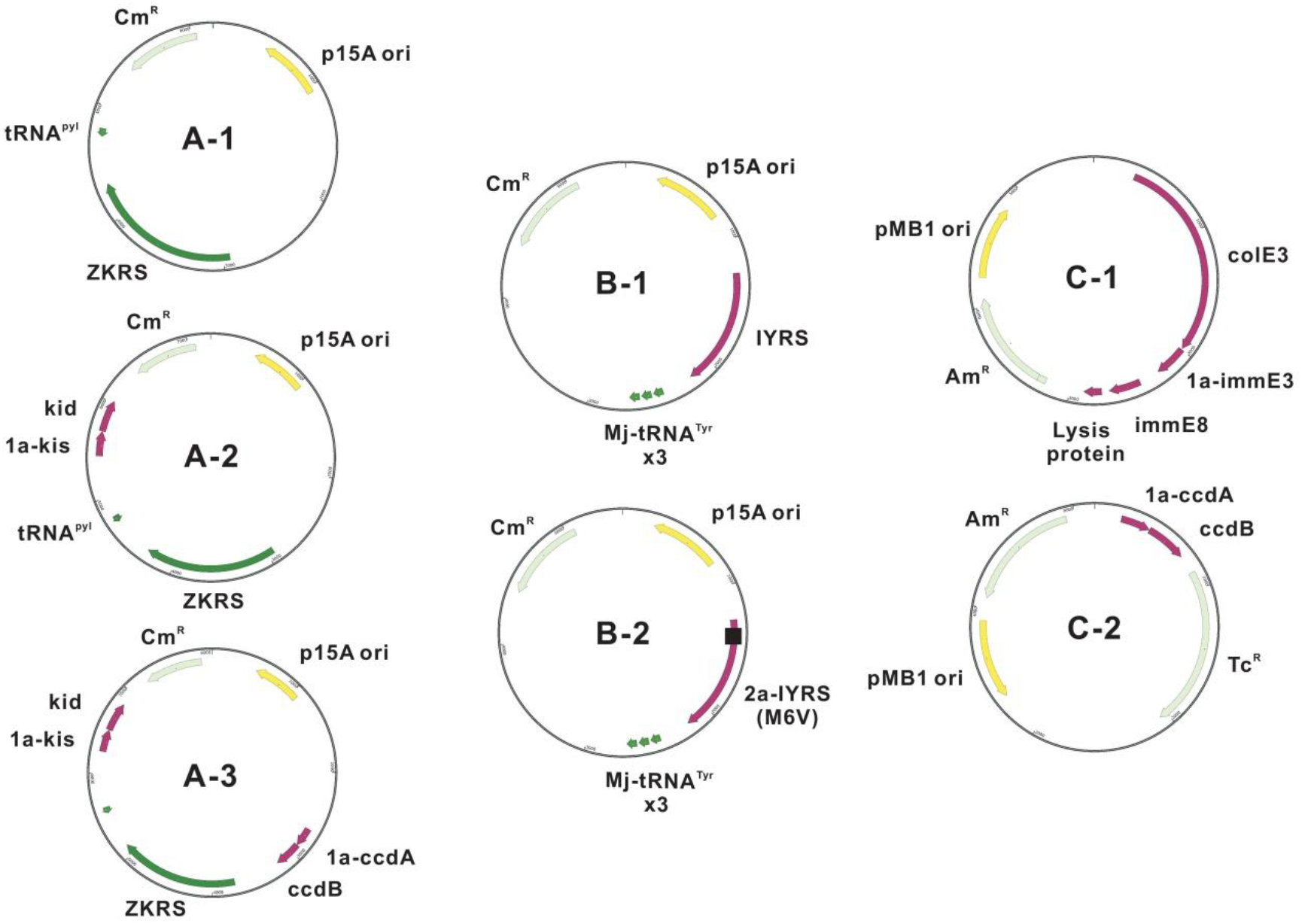
Plasmid maps. The plasmids used to construct the uAA-auxotrophic suicidal bacterial vaccines are shown. Plasmid A and B contain the p15A origin and the ZK or IY incorporation system consisting of a specific aminoacyl-tRNA synthetase and its cognate tRNA_CUA_. Although A-1 contains only the ZK incorporation system, A-2 and A-3 also carry *kid-kis* and both *kid-kis* and *ccdB-ccdA,* respectively. A TAG stop codon sequence, abbreviated as 1a, was inserted next to the translation start codon of antitoxin genes. Whereas B-1 maintains the intact IY incorporation system, B-2 contains a modified system to reduce leakage expression in the absence of IY using a positive-feedback loop, 2a-IYRS(M6V) (42). C-1 and C-2 carry the toxin-antitoxin gene *colE3-immE3* and *ccdB-ccdA* with the TAG insertion, respectively. Plasmid C is a pMB1 origin plasmid which can be co-transformed with plasmid A or B.

**Supplementary Figure 2.**
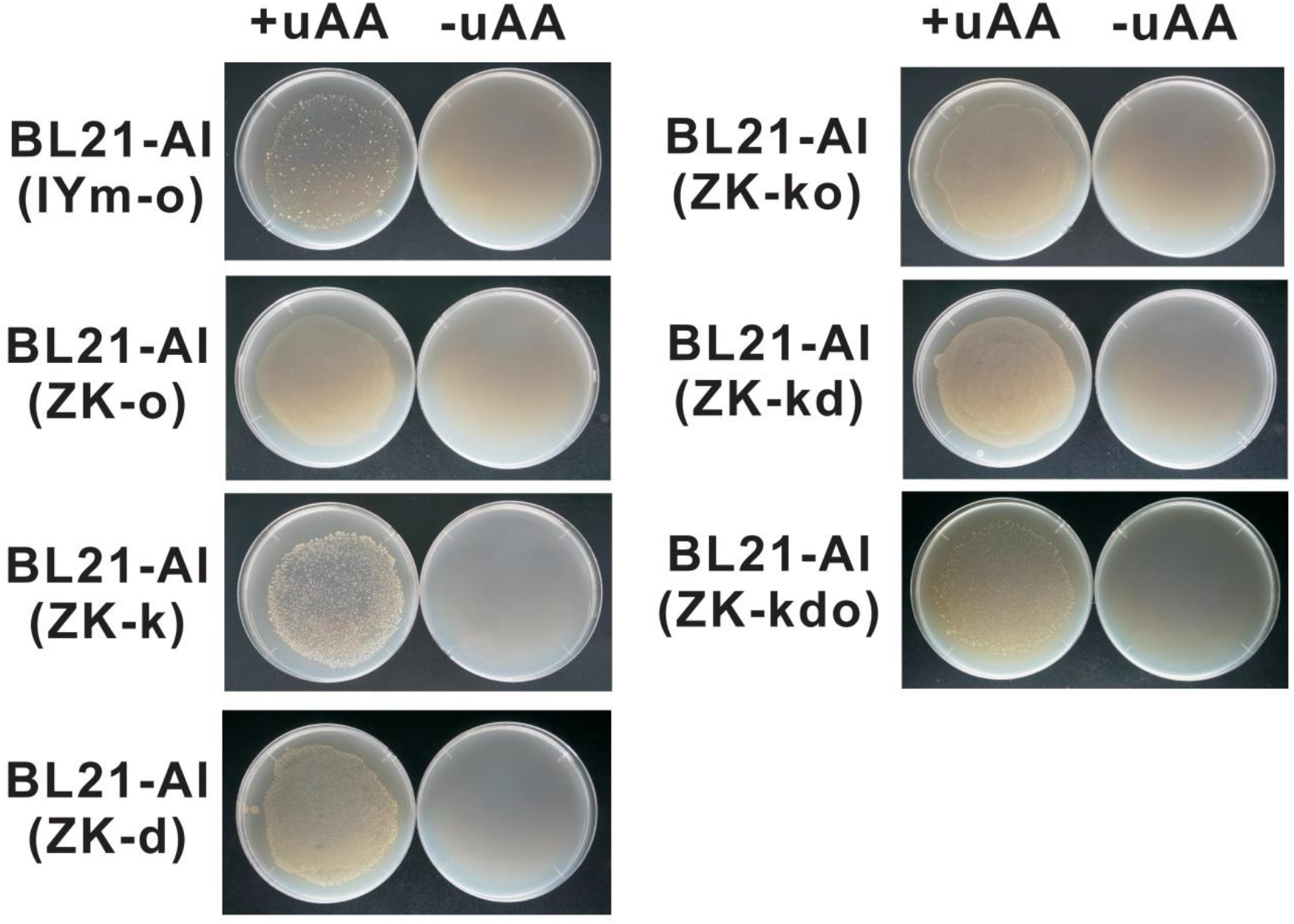
The uAA-auxotrophic *E. coli* BL21-AI laboratory strains. Various combination of plasmids shown in Supplementary Figure 1 were transfected into BL21-AI. Growth was evaluated in the presence and absence of 1 mM IY for BL21-AI(IYm-o) or 3 mM ZK for other strains.

**Supplementary Figure 3.**
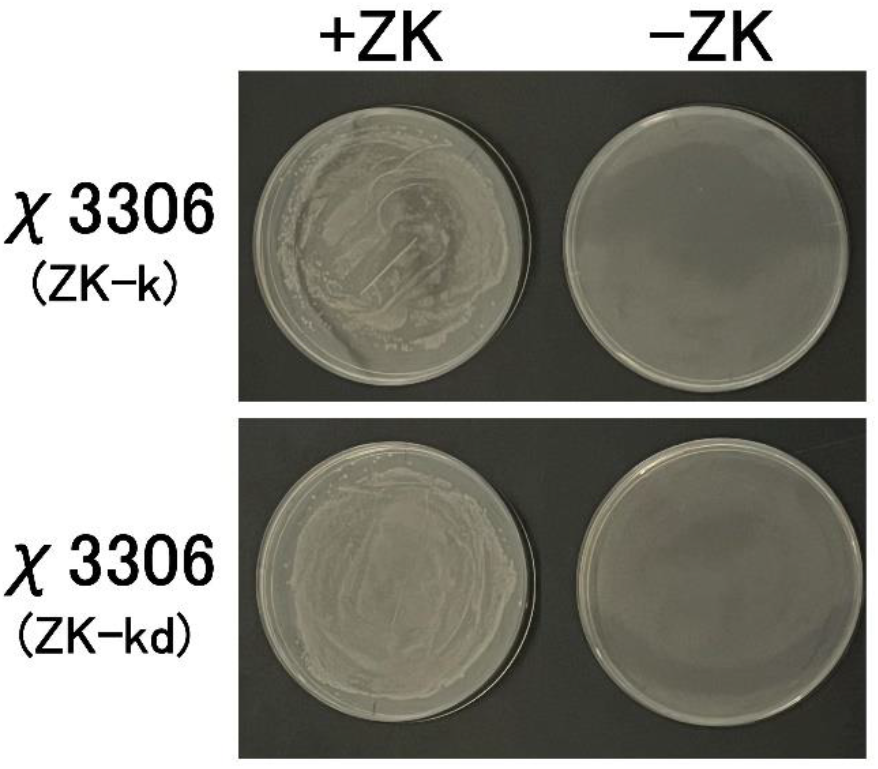
The uAA-auxotrophic *S. enterica* strains. *S. enterica* χ3306 (ZK-k) and χ3306 (ZK-kd) harbor plasmids A-2 and A-3, respectively. Growth was evaluated in the presence and absence of 3 mM ZK.

**Supplementary Figure 4.**
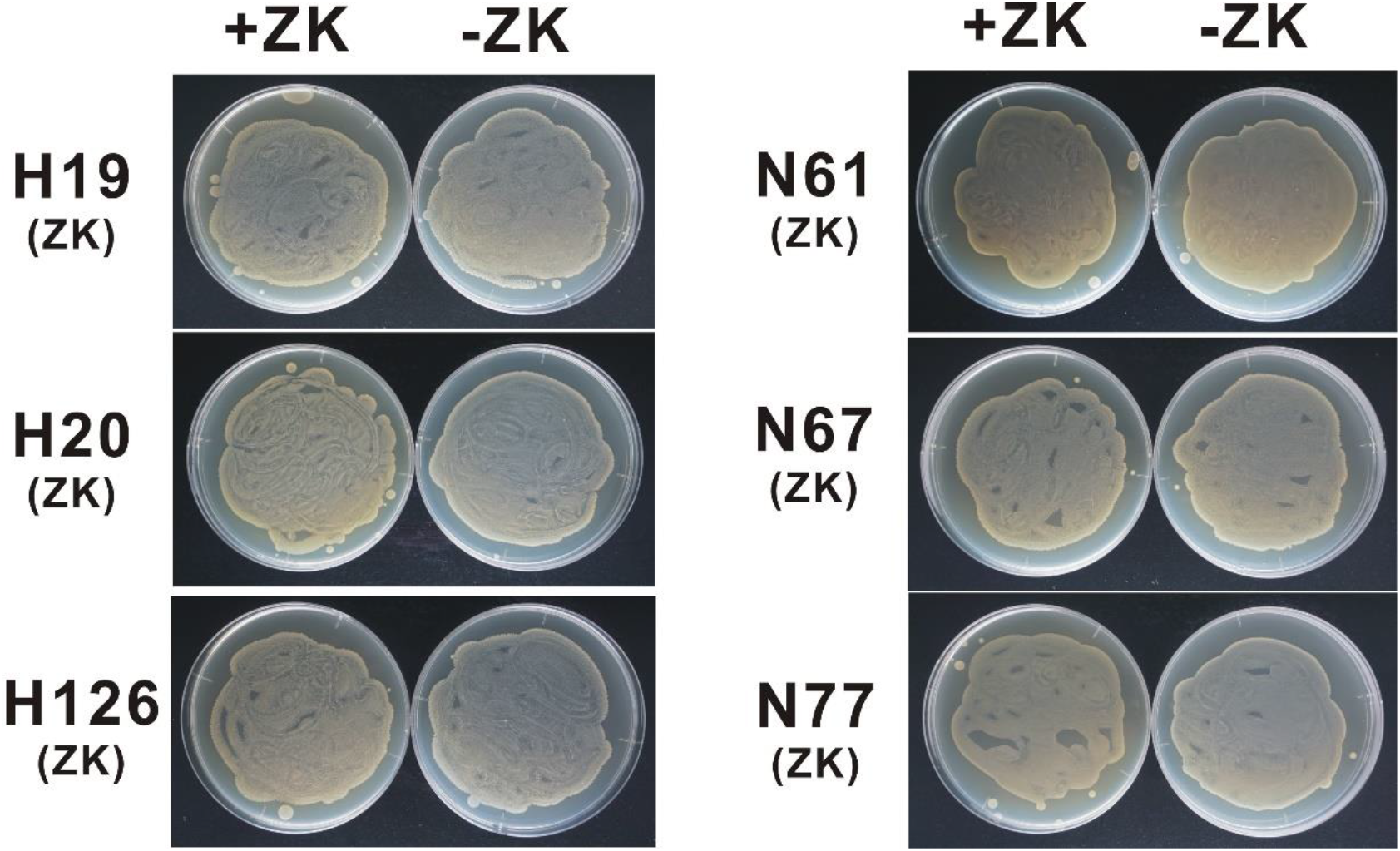
*E. coli* clinical isolates were tolerant for ZK-incorporation. Plasmid A-1 was transfected. Growth was tested in the presence and absence of 3 mM ZK.

**Supplementary Figure 5.**
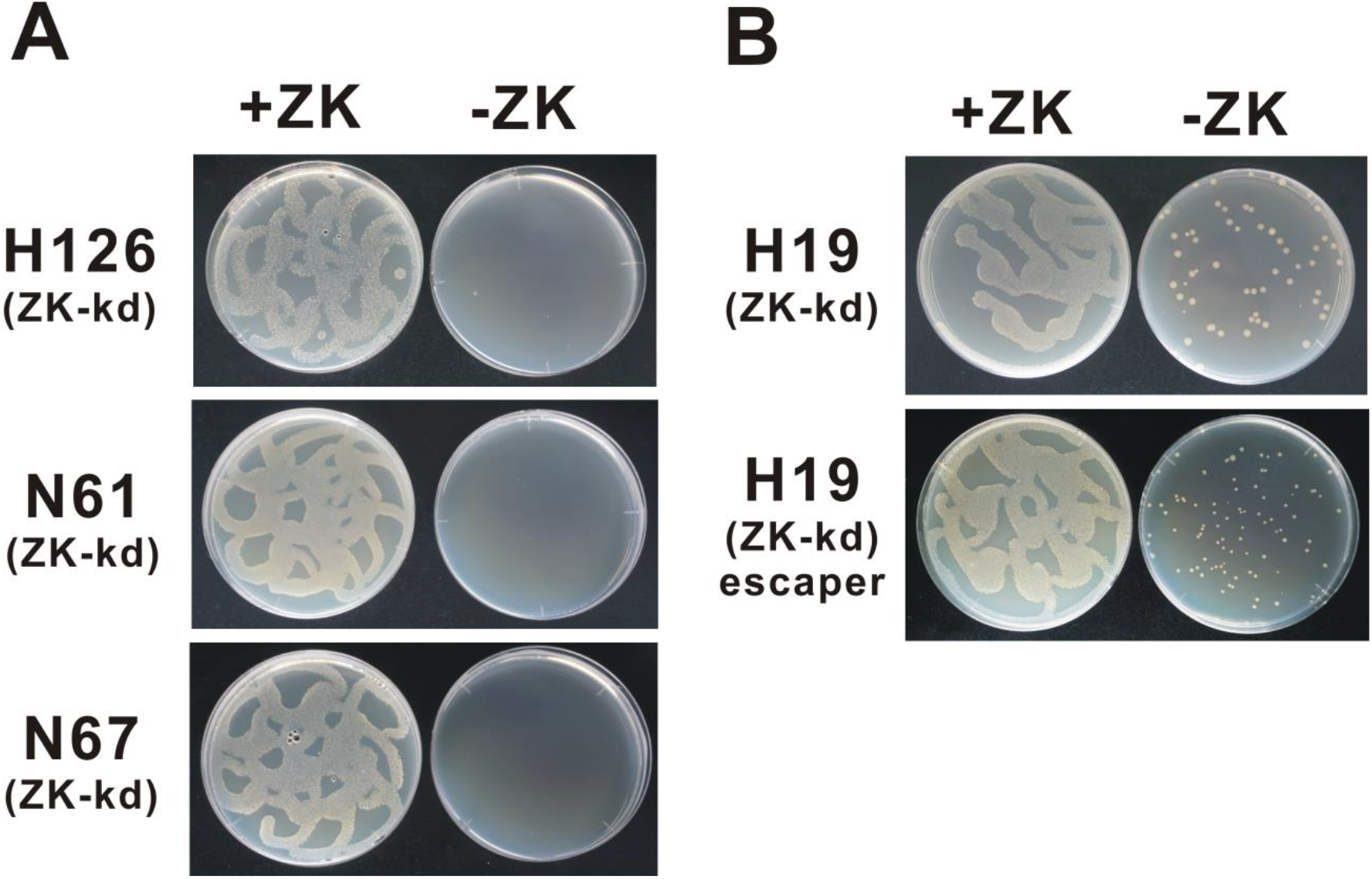
Transformation of the ZK-regulated *kid-kis* and *ccdB-ccdA* toxin-antitoxin systems into *E. coli* clinical isolates. Plasmid A-3 was transfected into *E. coli* clinical isolates. Transformants were successfully generated for 4 strains, H19, H126, N61 and N67. Colony formation was evaluated in the presence and absence of 3 mM ZK. Uneven colony distribution is an artificial phenomenon caused by uneven inoculation. See also Materials and Methods. (A) H126, N61 and N67. (B) H-19. Note a higher emergence of escapers even in the absence of ZK. Top panel, the first generation after the plasmid A-3 transformation. Bottom panel, a recultured escaper isolated from the first generation.

**Supplementary Figure 6.**
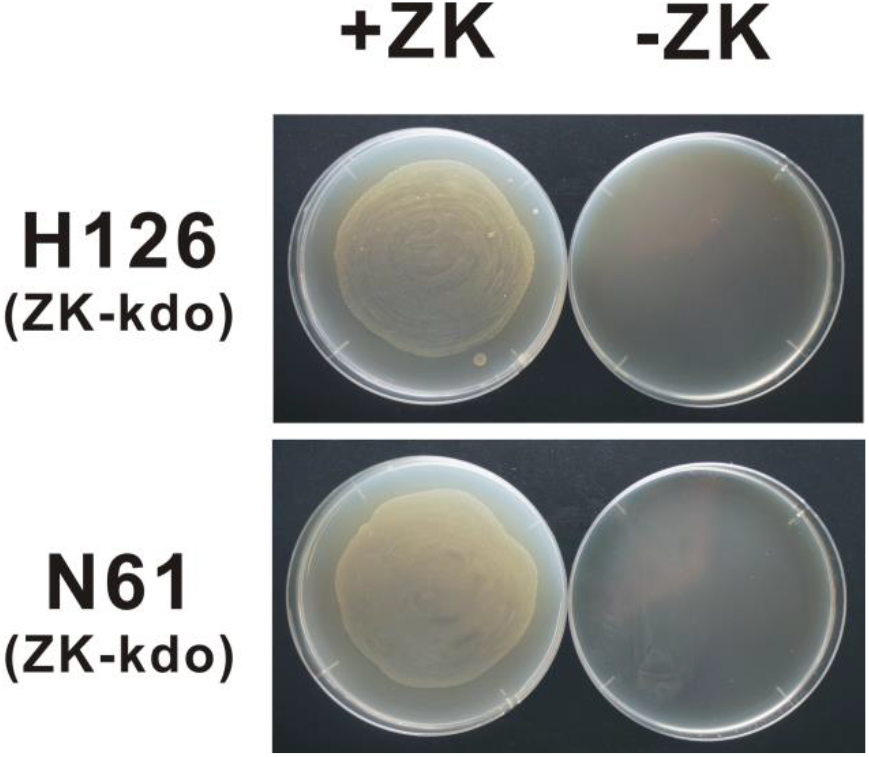
Additional transformation of the ZK-regulated *colE3-immE3* toxin-antitoxin system. Plasmid C-1 was additionally transformed into the resulting strains shown in Supplementary Figure 5. Growth was evaluated in the presence and absence of 3 mM ZK.

**Supplementary Table 1.**
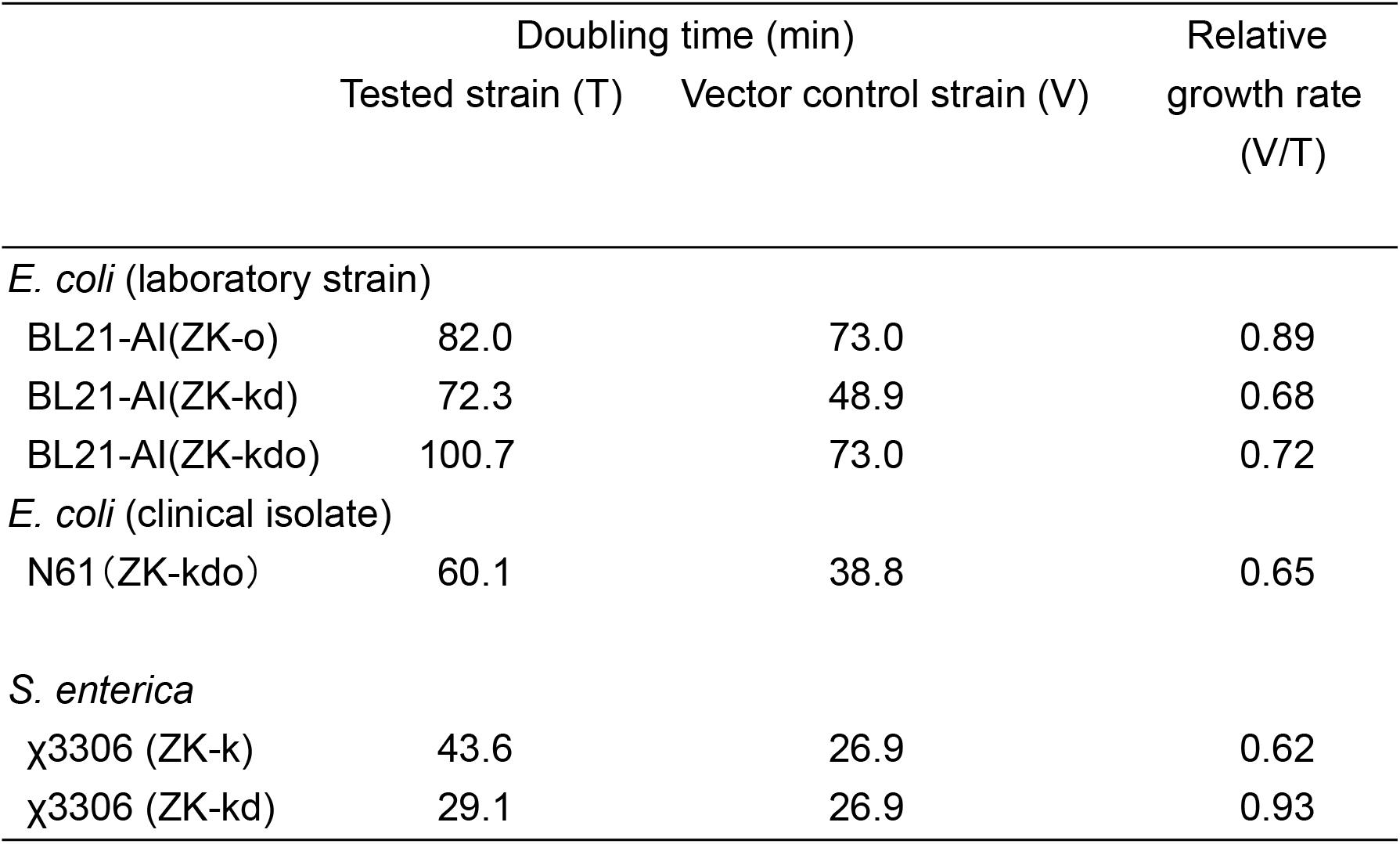
Growth rates. Growth curves were determined for some ZK-auxotrophic strains and their vector control strains. Doubling times were calculated during the logarithmic growth phase. Growth rates were shown as a relative value (vector control = 1).

**Supplementary Table 2.** Disease score.

| Disease Score | Characterization | Clinical signs |
| --- | --- | --- |
| 0 | No clinical signs | - |
| 1 | Mild clinical signs<sup>a</sup> | Ruffled fur |
| 2 | Moderate clinical signs | Ruffled fur plus, lethargy, hunched posture, decreased activity |
| 3 | Severe clinical signs | Paresis, paralysis, tremor, shivers, ataxia, rigidity, coma |
| 4 | Death | - |

<sup>a</sup> Mice that exhibited only mild clinical signs, showed no further signs of disease, and recovered until the end of the experiment were not considered ill.

## References

1. A. S. Fauci, It Ain’t Over Till It’s Over…but It’s Never Over—Emerging and Reemerging Infectious Diseases. N. Engl. J. Med. 387, 2009–2011 (2022).

2. A. Findlater, I. I. Bogoch, Human mobility and the global spread of infectious diseases: a focus on air travel. Trends Parasitol. 34, 772–783 (2018)..

3. B. D. Dalziel, S. Kissler, J. R. Gog, C. Viboud, O. N. Bjørnstad, C. J. E. Metcalf, B. T. Grenfell, Urbanization and humidity shape the intensity of influenza epidemics in U.S. cities. Science 362, 75–79 (2018).

4. S. Rauch, E. Jasny, K. E. Schmidt, B. Petsch, New vaccine technologies to combat outbreak situations. Front. Immunol. 9, 1963 (2018)..

5. E. S. Pronker, T. C. Weenen, H. Commandeur, E. H. Claassen, A. D. Osterhaus, Risk in vaccine research and development quantified. PLoS One 8, e57755 (2013).

6. B. Shanmugaraj, A. Malla, W. Phoolcharoen, Emergence of novel coronavirus 2019-nCoV: need for rapid vaccine and biologics development. Pathogens, 9, 148 (2020).

7. M. Vouga, G. Greub, Emerging bacterial pathogens: the past and beyond. Clin. Microbiol. Infect. 22, 12–21 (2016).

8. M. Brisse, S. M. Vrba, N. Kirk, Y. Liang, H. Ly, Emerging Concepts and Technologies in Vaccine Development. Front. Immunol. 11, 583077 (2020).

9. S. Black, D. E. Bloom, D. C. Kaslow, S. Pecetta, R. Rappuoli, Transforming vaccine development. Semin. Immunol. 50, 101413 (2020)

10. L. Pasteur, C. E. Chamberland, E. Roux, Sur la vaccination charbonneuse. C. R. Acad. Sci. 92, 1378–1383 (1881).

11. S. Plotkin, History of vaccination. Proc. Natl. Acad. Sci. U.S.A. 111, 12283–12287 (2014).

12. J. E. Galen, R. Curtiss III, The delicate balance in genetically engineering live vaccines. Vaccine 32, 4376–4385 (2014).

13. Y. Kato, An engineered bacterium auxotrophic for an unnatural amino acid: a novel biological containment system. PeerJ 3, e1247 (2015).

14. C. F. Shuster, R. Bertram, Toxin–antitoxin systems are ubiquitous and versatile modulators of prokaryotic cell fate. FEMS Microbiol. Lett. 340, 73–85 (2013).

15. D. de la Torre, J. W. Chin, Reprogramming the genetic code. Nat. Rev. Genet. 22, 169–184 (2021).

16. M. Minaba, Y. Kato, (2014). High-yield, zero-leakage expression system with a translational switch using site-specific unnatural amino acid incorporation. Appl. Environ. Microbiol. 80, 1718–1725 (2014).

17. Y. Kato, Translational Control using an Expanded Genetic Code. Int. J. Mol. Sci. 20, 887 (2019).

18. G. Dietrich, et al., Delivery of antigen-encoding plasmid DNA into the cytosol of macrophages by attenuated suicide *Listeria monocytogenes*. Nat. Biotechnol. 16, 181–185 (1998).

19. W. Kong, et al., Regulated programmed lysis of recombinant *Salmonella* in host tissues to release protective antigens and confer biological containment. Proc. Natl. Acad. Sci. U.S.A. 105, 9361–9366 (2008).

20. Z. Hindle, et al. Characterization of *Salmonella enterica* derivatives harboring defined *aroC* and *Salmonella* pathogenicity island 2 type III secretion system (*ssaV*) mutations by immunization of healthy volunteers. Infect. Immun. 70, 3457–3467 (2002).

21. S. M. Tennant, M. M. Levine, Live attenuated vaccines for invasive *Salmonella* infections. Vaccine 33, C36–C41 (2015).

22. L. Pöyhönen, et al. Life-threatening infections due to live-attenuated vaccines: early manifestations of inborn errors of immunity. J Clin. Immunol. 39, 376–390 (2019).

23. A. J. Cann, et al. Reversion to neurovirulence of the live-attenuated Sabin type 3 oral pollovirus vaccine. Nucleic Acids Res. 12, 7787–4492 (1984).

24. C. M. Whitford, et al. Auxotrophy to Xeno-DNA: an exploration of combinatorial mechanisms for a high-fidelity biosafety system for synthetic biology applications. J. Biol. Eng. 12, 13 (2018).

25. L. Torres, A. Krüger, E. Csibra, E. Gianni, V. B. Pinheiro, V. B., Synthetic biology approaches to biological containment: pre-emptively tackling potentioal risks. Essays Biochem. 60, 393–410 (2016).

26. F. Stirling, P. A. Silver, P. A., Controlling the implementation of transgenic microbes: Are we ready for what synthetic biology has to offer? Mol. Cell 78, 614–623 (2020).

27. J. W. Lee, C. T. Y. Chan, S. Slomovic, J. J. Collins, Next-generation biocontainment systems for engineered organisms. Nat. Chem. Biol. 14, 530–537 (2018).

28. H. J. O. Ramos, M. G. Yates, F. O. Pedrosa, E. M. Souza EM., Strategies for bacterial tagging and gene expression in plant-host colonization studies. Soil Biol. Biochem. 43, 1626–1638 (2011).

29. O. Wright, M. Delmans, G.-B. Stan, T. Ellis, 2014. GeneGuard: a modular plasmid system designed for biosafety. ACS Synth. Biol. 4, 307–316 (2014).

30. Y. Kato, A strategy for addicting transgene-free bacteria to synthetic modified metabolites. Front. Microbiol. 14, 1086094 (2023)

31. T. Dandeka, A. Fieselmann, J. Popp, M. Hensel, M, *Salmonella enterica*: a surprisingly well-adapted intracellular lifestyle. Front. Microbiol. 3, 164 (2012).

32. J. Kaper, J. Nataro, H. Mobley, Pathogenic *Escherichia coli*. Nat. Rev. Microbiol. 2, 123–140 (2004).

33. WHO, “WHO Product Development for Vaccines Advisory Committee (PDVAC). Update on Development of Enterotoxigenic E. Coli (ETEC) Vaccines.” (2020). Available at: https://cdn.who.int/media/docs/default-source/immunization/pdvac/2020/pdvac_etec-vaccines_18-june_presentations_reduced.pdf?sfvrsn=930ceb78_8

34. S. M. Baliban, Y.-J. Lu, R. Malley, Overview of the nontyphoidal and paratyphoidal *Salmonella* vaccine pipeline: current status and future prospects, Clin. Infect. Dis. 71, S151–S154 (2020).

35. M. Fukushima, K. Kakinuma, R. Kawaguchi, Phylogenetic analysis of *Salmonella*, *Shigella*, and *Escherichia coli* strains on the basis of the *gyrB* gene sequence. J. Clin. Microbiol. 40, 2279–2785 (2002).

36. C. M. VanDrisse, J. C. Escalante-Semerena, New high-cloning-efficiency vectors for complementation studies and recombinant protein overproduction in *Escherichia coli* and *Salmonella enterica*. Plasmid 86, 1–6 (2016).

37. N. Bhawsinghka, K. F. Glenn, R. M. Schaaper, Complete genome sequence of *Escherichia coli* BL21-AI. Microbiol. Resour. Announc. 9, e00009–20 (2020).

38. H. Masaki, T. Ohta, Colicin E3 and its immunity genes. J. Mol. Biol. 182, 217–227 (1985).

39. NIH (2019). “NIH Guidelines for Research Involving Recombinant or Synthetic Nucleic Acid Molecules.” (2019). Available at: https://osp.od.nih.gov/ufaq-category/nih-guidelines-for-research-involving-recombinant-or-synthetic-nucleic-acidmolecules/

40. Y. Kato, Tunable translational control using site-specific unnatural amino acid incorporation in *Escherichia coli*. PeerJ 3, e904 (2015).

41. K. Sakamoto, et al., Genetic encoding of 3-iodo-_L_-tyrosine in *Escherichia coli* for single-wavelength anomalous dispersion phasing in protein crystallography. Structure, 17, 335–344 (2009).

42. Y. Kato, Tight translational control using site-specific unnatural amino acid incorporation with positive feedback gene circuits. ACS Synth. Biol. 7, 1956–1963 (2018).

43. T. Yanagisawa, et al., Multistep engineering of pyrrolysyl-tRNA synthetase to genetically encode *N*^ɛ^-(o-azidobenzyloxycarbonyl) lysine for site-specific protein modification. Chem. Biol. 15, 1187–1197 (2008).

44. E. Diago-Navarro, et al., (2010). *parD* toxin–antitoxin system of plasmid R1–basic contributions, biotechnological applications and relationships with closely-related toxin–antitoxin systems. FEBS J. 277, 3097–3117 (2010).

45. E. M. Bahassi, et al., Interactions of CcdB with DNA gyrase: inactivation of GyrA, poisoning of the gyrase-DNA complex, and the antidote action of CcdA. J. Biol. Chem. 274, 10936–10944 (1999).

46. B. Torres, S. Jaenecke, K. N. Timmis, J. L. García, E. Díaz, (2003). A dual lethal system to enhance containment of recombinant micro-organisms. Microbiology 149, 3595–3601 (2003).

47. R. R. Gallagher, J. R. Patel, A. L. Interiano, A. J. Rovner, F. J. Isaacs, Multilayered genetic safeguards limit growth of microorganisms to defined environments. Nucleic Acids Res. 43, 1945–1954 (2015).

48. P. A. Gulig, R. Curtiss III, Plasmid-associated virulence of Salmonella typhimurium. Infect. Immun. 55, 2891–2901 (1987).

49. K. Sharun, et al., Advances in therapeutic and managemental approaches of bovine mastitis: a comprehensive review, Veterinary Quarterly 41, 107–136 (2021).

50. H. Tian, et al., Screening system for orthogonal suppressor tRNAs based on the species-specific toxicity of suppressor tRNAs. Biochimie, 95, 881–888 (2013).

51. D. Jurėnas, N. Fraikin, F. Goormaghtigh, L. Van Melderen, Biology and evolution of bacterial toxin–antitoxin systems. Nat. Rev. Microbiol. 20, 335–350 (2022).

52. T. Mukai, et al., Codon reassignment in the *Escherichia coli* genetic code. Nucleic Acids Res. 38, 8188–8195 (2010).

53. D. B. Johnson, et al., RF1 knockout allows ribosomal incorporation of unnatural amino acids at multiple sites. Nat. Chem. Biol. 7, 779–786 (2011).

54. M. M. Gibani, C. Britto, A. J. Pollard, Typhoid and paratyphoid fever: a call to action. Curr. Opin. Infect. Dis. 31, 440–448 (2018)

55. C. V. Pulford, et al., Stepwise evolution of *Salmonella Typhimurium* ST313 causing bloodstream infection in Africa. Nat. Microbiol. 6, 327–338 (2021)

56. F. Vangroenweghe, L. Duchateau, C. Burvenich, J-5 *Escherichia coli* vaccination does not influence severity of an *Escherichia coli* intramammary challenge in primiparous cows. J. Dairy Sci. 103, 6692–6697 (2020)..

57. R. Turk, R., et al., Milk and serum proteomes in subclinical and clinical mastitis in Simmental cows. J. Proteomics, 244, 104277 (2021).

58. G. Yang, G., et al., Antibacterial and immunomodulatory effects of Pheromonicin-NM on *Escherichia coli*-challenged bovine mammary epithelial cells. Int. Immunopharmacol. 84, 106569 (2020).

59. M. Toquet, M., Á. Gómez-Martín, E. Bataller, Review of the bacterial composition of healthy milk, mastitis milk and colostrum in small ruminants. Res. Vet. Sci. 140, 1–5 (2021).

60. R. J. Goldstone, S. Harris, D. G. Smith, Genomic content typifying a prevalent clade of bovine mastitis-associated *Escherichia coli*. Sci. Rep. 6, 30115 (2016).

61. A. X. Tran, C. Whitfield, “Lipopolysaccharides (endotoxins)” in Encychlopedia of Microbiology (Third *Edition)*, M. Schaechter Ed. (Elsevier, 2009), pp.513–528.

62. Y. Xu, M. Zhou, C. Zhu, W. Ma, Construction of recombinant strain of low toxic lipopolysaccharide-producing *Escherichia coli* J5 gene. Jiangsu Agric. Sci. 48, 60–64 (2020)

63. P. Rainard, M. Repérant-Ferter, C. Gitton, P. Germon, Shielding Effect of *Escherichia coli* O-antigen polysaccharide on J5-induced cross-reactive antibodies. mSphere, 6, e01227–20 (2021).

64. N. Zaatout, An overview on mastitis-associated *Escherichia coli*: Pathogenicity, host immunity and the use of alternative therapies. Microbiol. Res. 256, 126960 (2022).

65. R. H. Gregersen, H. Christensen, C. Ewers, M. Bisgaard, (2010). Impact of *Escherichia coli* vaccine on parent stock mortality, first week mortality of broilers and population diversity of *E. coli* in vaccinated flocks. Avian Pathol. 39, 287–295 (2010).

66. L. Li, I. Thøfner, J. P. Christensen, T. Ronco, K. Pedersen, R. H. Olsen, Evaluation of the efficacy of an autogenous *Escherichia coli* vaccine in broiler breeders. Avian Pathol. 46, 300–308 (2017).

67. R. Curtiss, Bacterial infectious disease control by vaccine development. J. Clin. Invest. 110, 1061–1066 (2002).

68. S. E. Luria, M. Delbrück, M., Mutations of bacteria from virus sensitivity to virus resistance. Genetics, 28, 491 (1943).

69. B. M. Hall, C. X. Ma, P. Liang, K. K. Singh, Fluctuation AnaLysis CalculatOR: a web tool for the determination of mutation rate using Luria–Delbrück fluctuation analysis. Bioinformatics 25, 1564–1565 (2009).

70. M. Eguchi, Y. Kikuchi, Binding of Salmonella-specific antibody facilitates specific T cell responses via augmentation of bacterial uptake and induction of apoptosis in macrophages. J. Infect. Dis. 201, 62–70 (2010)

71. S. D. Aribam, et al., Specific monoclonal antibody overcomes the *Salmonella enterica* serovar Typhimurium’s adaptive mechanisms of intramacrophage survival and replication. PLoS One 11, e0151352 (2016)

72. H. Matsui, S. Fukiya, C. Kodama-Akaboshi, M. Eguchi, T. Yamamoto, Mouse models for assessing the cross-protective efficacy of oral non-typhoidal *Salmonella* vaccine candidates harbouring in-frame deletions of the ATP-dependent protease *lon* and other genes. J. Med. Microbiol. 64(Pt 3), 295–302 (2015)

73. C. C. Toledo, et al., Shiga toxin-mediated disease in MyD88-deficient mice infected with *Escherichia coli* O157: H7. Am. J. Pathol. 173, 1428–1439 (2008).

